# The pheromone and pheromone receptor mating-type locus is involved in controlling uniparental mitochondrial inheritance in *Cryptococcus*

**DOI:** 10.1101/813998

**Authors:** Sheng Sun, Ci Fu, Giuseppe Ianiri, Joseph Heitman

## Abstract

Mitochondria are inherited uniparentally during sexual reproduction in the majority of eukaryotic species studied, including humans, mice, nematodes, as well as many fungal species. Mitochondrial uniparental inheritance (mito-UPI) could be beneficial in that it avoids possible genetic conflicts between organelles with different genetic backgrounds, as recently shown in mice; and it could prevent the spread of selfish genetic elements in the mitochondrial genome. Despite the prevalence of observed mito-UPI, the underlying mechanisms and the genes involved in controlling this non-mendelian inheritance are poorly understood in many species. In *Cryptococcus neoformans*, a human pathogenic basidiomyceteous fungus, mating types (*MAT*α and *MAT***a**) are defined by alternate alleles at the single *MAT* locus that evolved from fusion of the two *MAT* loci (*P/R* encoding pheromones and pheromone receptors, *HD* encoding homeodomain transcription factors) that are the ancestral state in the basidiomycota. Mitochondria are inherited uniparentally from the *MAT***a** parent in *C. neoformans* and this requires the *SXI1*α and *SXI2***a** HD factors encoded by *MAT*. However, there is evidence additional genes contribute to control of mito-UPI in *Cryptococcus*. Here we show that in *Cryptococcus amylolentus*, a sibling species of *C. neoformans* with unlinked *P/R* and *HD MAT* loci, mitochondrial uniparental inheritance is controlled by the *P/R* locus, and is independent of the *HD* locus. Consistently, by replacing the *MAT*α alleles of the pheromones (*MF*) and pheromone receptor (*STE3*) with the *MAT***a** alleles, we show that these *P/R* locus defining genes indeed affect mito-UPI in *C. neoformans* during sexual reproduction. Additionally, we show that during early stages of *C. neoformans* sexual reproduction, conjugation tubes are always produced by the *MAT*α cells, resulting in unidirectional migration of the *MAT*α nucleus into the *MAT***a** cell during zygote formation. This process is controlled by the *P/R* locus and could serve to physically restrict movement of *MAT*α mitochondria in the zygotes, and thereby contribute to mito-UPI. We propose a model in which both physical and genetic mechanisms function in concert to prevent the coexistence of mitochondria from the two parents in the zygote and subsequently in the meiotic progeny, thus ensuring mito-UPI in pathogenic *Cryptococcus*, as well as in closely related non-pathogenic species. The implications of these findings are discussed in the context of the evolution of mito-UPI in fungi and other more diverse eukaryotes.

## INTRODUCTION

Mitochondria are important eukaryotic organelles. In addition to providing cellular energy, they are involved in a variety of cellular processes, such as signaling, cellular differentiation, cell death, as well as control of the cell cycle and cell growth (MCBRIDE *et al.* 2006). Furthermore, they are implicated in several human diseases, such as mitochondrial disorders and cardiac dysfunction, and may play a role in aging (LESNEFSKY *et al.* 2001; GARDNER AND BOLES 2005). In pathogenic fungi, such as *Cryptococcus neoformans*, mitochondria play critical roles in virulence, survival under low oxygen conditions, and drug tolerance (INGAVALE *et al.* 2008; MA *et al.* 2009; MA AND MAY 2010; SHINGU-VAZQUEZ AND TRAVEN 2011; KRETSCHMER *et al.* 2012). Additionally, given effects of mitochondrial dysfunction on fungal drug tolerance and virulence, these organelles are potential antifungal drug development targets (SHINGU-VAZQUEZ AND TRAVEN 2011).

Unlike nuclear genes and genomes, the inheritance of mitochondrial (as well as chloroplast) genes and genomes does not follow Mendel’s laws. These organelle genomes are typically uniparentally inherited in the majority of species that have been studied. There are currently two hypotheses for the evolution of mitochondrial uniparental inheritance (UPI). First, mito-UPI evolved as a mechanism to restrict spread of mitochondrial selfish elements that enhance mitochondrial fitness to the detriment of their host. Second, mito-UPI evolved to avoid having two genetically different mitochondrial genomes in the zygote simultaneously. This prevents mitochondrial heteroplasmy and/or recombination, which are thought to either generate less fit genomes, or cause nuclear-mitochondrial/mitochondrial-mitochondrial incompatibilities. Although it is yet not known why mitochondrial heteroplasmy is deleterious, mice that inherit both parental mitochondrial genomes exhibit signs of cellular or organismal imbalance associated with a variety of phenotypes, including behavioral and cognitive abnormalities (LANE 2012; SHARPLEY *et al.* 2012). This could be the result of Muller-Dobzhansky type incompatibilities when one nuclear genome must work in concert with two distinct types of mitochondrial genomes (MAHESHWARI AND BARBASH 2011).

Among fungal species that have been examined, a majority exhibits uniparental mitochondrial inheritance, that is, the progeny all possess a mitochondrial genome inherited from one of the two mating parents. Examples of fungal uniparental mitochondrial inheritance include *Aspergillus nidulans, Neurospora crassa, Candida albicans, Coprinopsis cinerea, Agaricus bisporus, Cryptococcus neoformans*, and *Ustilago maydis* (XU 2005; GYAWALI AND LIN 2011; NI *et al.* 2011; SHAKYA AND IDNURM 2013; GOODENOUGH AND HEITMAN 2014). In contrast, in some fungal species such as *Saccharomyces cerevisiae* and *Schizosaccharomyces pombe*, inheritance of mitochondrial genomes is biparental. In these species, mating between isogametic sexual partners results in an equal contribution of organelles from the two gametes into the zygote, and the transient coexistence of two different mitochondria often results in recombination (EGAL *et al.* 1980; BIRKY 1996; BIRKY 2001). However, even in the cases of biparental inheritance, homoplasy is rapidly re-established after the initial heteroplasmic zygote, and each of the daughter cells possesses the mitochondrion of one genetic type, either from one of the two parents, or a recombinant of the two (BIRKY 1996; BERGER AND YAFFE 2000; BIRKY 2001; BARR *et al.* 2005).

Several mechanisms have been proposed to explain mito-UPI. In anisogametic species, such as animals and plants, the size differences between male and female gametes result in biased contributions of organelles in the zygotes, which could serve to restrict the transmission of mitochondria from the paternal parent. In addition, active mitochondrial marking and degradation during zygote formation have also been identified in several species (AL RAWI *et al.* 2011; LEVINE AND ELAZAR 2011; SATO AND SATO 2011; ZHOU *et al.* 2011; LUO *et al.* 2013). For example, ubiquitin marking followed by mitophagy of unmarked sperm mitochondria shortly after zygote formation has recently been implicated in mito-UPI in *Caenorhabditis elegans*.

Most fungi are isogametic and mating occurs between two gametes of the same or similar size, or between two compatible mycelia. Thus, unlike plants and animals, where size differences between the gametes contributes to unequal organelle contributions to the zygote, and subsequently facilitates uniparental organelle inheritance, in fungi, several different mechanisms have evolved that could actively avoid mitochondrial heteroplasy during sexual reproduction (WILSON AND XU 2012). For example, in some fungal species (e.g. *Coprinus cinereus*), mating between two compatible mycelia is achieved by unidirectional migration of nuclei while all of the cytoplasm including the organelles is left behind; thus, the mixing of two different mitochondria is avoided (HINTZ *et al.* 1988; MAY AND TAYLOR 1988).

In unicellular fungi where mating occurs between two isogametic mating partners, there is evidence that organelles from one parent are actively degraded, thus ensuring homoplasy in the zygote. One example is the plant pathogenic fungus *Ustilago maydis* (FEDLER *et al.* 2009). *U. maydis* is a basidiomycete and has a tetrapolar mating system, constituted by the *a* and *b* loci. The biallelic *a* locus (*a1* and *a2*) is involved in pheromone and pheromone receptor based cell recognition and fusion, while the multi-allelic *b* locus encodes the homeodomain transcription factors. In *U. maydis*, mitochondrial inheritance is governed by the *a2*-specific genes *LGA2* and *RGA2*. The Lga2 and Rga2 proteins both localize to mitochondria, and Lga2 interferes with mitochondrial dynamics and fusion (BORTFELD *et al.* 2004; MAHLERT *et al.* 2009). Evidence also supports an active Lga2- and Rga2-mediated selective mitochondrial elimination process. Specifically, Lga2 is the destroyer, produced by the a2 locus and needed to destroy the mitochondria from the a1 parent, while Rga2 is the protector, produced by the a2 locus and needed to protect the a2 mitochondria from mitophagic destruction. However, the fact that deletion of *RGA2* reverses the inheritance in favor of the *a1*-type mitochondria (rather than a biparental pattern) indicates that there might also exist another *RGA2-*independent mechanism that is involved in the control of mitochondrial inheritance (FEDLER *et al.* 2009), although it has been reported that the mitophagy related gene *ATG11* is not required for mito-UPI in *U. maydis* (WAGNER-VOGEL *et al.* 2015).

*Cryptococcus neoformans* is a human pathogenic basidiomycete fungus that was classified into two varieties: var. *grubii* (serotype A) and var. *neoformans* (serotype D), that are now recognized as distinct species, *C. neoformans* (serotype A) and *C. deneoformans* (serotype D) (HAGEN *et al.* 2015). *C. neoformans* has a defined bipolar mating system with two mating types (α and **a**), which are defined by the alleles present at the mating type locus (*MAT*) (KWON-CHUNG 1975). Compared to other basidiomycetes, the *C. neoformans MAT* locus is unusually large, spanning more than 100 kb and containing more than 20 genes. Studies have shown that the *MAT* locus in *C. neoformans* is a fusion product of the ancestral *P/R* and *HD* loci, and thus includes genes encoding both homeodomain transcription factors (*HD*), as well as pheromones and pheromone receptors (*P/R*) (LOFTUS *et al.* 2005; HSUEH *et al.* 2011; SUN *et al.* 2017; PASSER *et al.* 2019). Mating in *C. neoformans* typically occurs between strains of opposite mating types (α and **a**), and mitochondrial inheritance during these opposite-sex matings is uniparental from the **a** parent (XU *et al.* 2000; YAN AND XU 2003).

Opposite-sex mating can also occur between strains of different species (*C. neoformans* and *C. deneoformans*), and mito-UPI from the *MAT***a** parent remains intact during these inter-species opposite-sex matings (YAN *et al.* 2007a). Additionally, mating in *C. neoformans* can also occur between two α strains, and mitochondrial inheritance during these same-sex matings has been shown to be biparental from both parents (YAN *et al.* 2007a). Mito-UPI in *C. neoformans* is established rapidly after zygote formation, and several genes play important roles in this process. For example, the two homeodomain transcription factors, *SXI1*α (in *MAT*α) and *SXI2***a** (in *MAT***a**), are both required to ensure mito-UPI, and deletion of either gene results in leakage of mitochondria from the *MAT*α parent to the zygote (YAN *et al.* 2004; YAN *et al.* 2007a). Interestingly, when the *SXI1*α and *SXI2***a** genes were exchanged between mating types, progeny still inherited mitochondria from the *MAT***a** parent, which now has a *sxi2***a**Δ deletion and a transgenic copy of *SXI1*α, indicating that other genes in the *MAT* locus are required for mito-UPI (HSUEH *et al.* 2008). Recently, Mat2, which is a pheromone activated transcription factor not encoded by *MAT*, was shown to be involved in mito-UPI in *C. neoformans* (GYAWALI AND LIN 2013). However, it is still not known what gene(s) in the *C. neoformans MAT* locus is the master regulator that initiates mito-UPI. Additionally, it is yet to be determined whether mito-UPI in *C. neoformans* also involves active mitochondrial marking and subsequent degradation via mitophagy or other processes, similar to mechanisms operating in *C. elegans* and *U. maydis* (FEDLER *et al.* 2009; AL RAWI *et al.* 2011; LEVINE AND ELAZAR 2011; SATO AND SATO 2011; ZHOU *et al.* 2011; LUO *et al.* 2013).

*Cryptococcus amylolentus*, together with *Cryptococcus floricola* and *Cryptococcus wingfieldii*, are the most closely related known sibling species of the *C. neoformans*/*C. gattii* pathogenic species complex (FINDLEY *et al.* 2009; PASSER *et al.* 2019). *C. amylolentus* has a tetrapolar mating system with the two mating type loci (*A* and *B*) located on different chromosomes. The A (*P/R*) locus is ∼100 kb in size and encodes the pheromones and pheromone receptors (and others), while the B (*HD*) locus is less than 10 kb in size and contains both *SXI1*α and *SXI2***a** homologs (FINDLEY *et al.* 2012). During *C. amylolentus* sexual reproduction, mitochondrial inheritance is uniparental and from the A2B2 parent (FINDLEY *et al.* 2012). However, due to a lack of proper mitochondrial-nuclear marker combinations, we were not able to further dissect which *MAT* locus, A or B or both, was responsible for mito-UPI in *C. amylolentus*.

In this study, we first obtained *C. amylolentus* isolates that have all eight possible mating type-mitochondrial combinations (*HD* – *P/R* – Mito). By analyzing mitochondrial inheritance in all possible pairwise crosses, we determined that mito-UPI in *C. amylolentus* is controlled by the *P/R* locus that encodes the pheromones and pheromone receptors, and is independent of the allele present at the *HD* locus. Additionally, we showed that in *C. neoformans*, the defining genes of the *P/R* locus in basidiomycetes (the pheromone and pheromone receptor genes) are indeed influencing mitochondrial inheritance during sexual reproduction. We also observed unidirectional conjugation tube formation from *MAT*α cells, a process controlled by the *P/R* locus, and subsequent polarized hyphal formation from the zygote, which together could act as physical barriers that prevent mitochondria of *MAT*α cells from spreading into the hyphae produced by the polarized zygote. Furthermore, we showed that deleting the *CRG1* gene, which encodes a regulator of G-protein signaling (RGS) that negatively regulates the pheromone-signaling cascade and cAMP pathways during sexual reproduction, resulted in *MAT*α and *MAT***a** strains with elevated responses to pheromones during mating. However, this enhanced response to pheromone is significantly more pronounced in the *MAT*α strains. Interestingly, progeny dissected from the *crg1*Δ bilateral crosses showed increased levels of mitochondrial leakage from the *MAT*α cells, consistent with the dynamics of pheromone sensing and the initial stages of sexual development playing an important role in mito-UPI. Taken together, we propose an integrated model that involves both physical and genetic mechanisms operating coordinately to enforce mito-UPI in *C. neoformans*, which is consistent with previous studies, and could serve as a foundation for future research on mitochondrial inheritance in *C. neoformans*, as well as in closely related basidiomycetes and beyond.

## MATERIALS AND METHODS

### Strains employed in this study

Strains used in this study, as well as their genotypes, are listed in Table 1. All strains were maintained in -80°C frozen stocks in 15% glycerol and subcultured from freezer stocks to YPD solid medium for study. Genotypic and phenotypic markers for the two laboratory constructed *C. neoformans* strains are listed in Table 1, as well as in previous studies where these strains were originally published (HSUEH *et al.* 2008; STANTON *et al.* 2010).

**Table 1.**
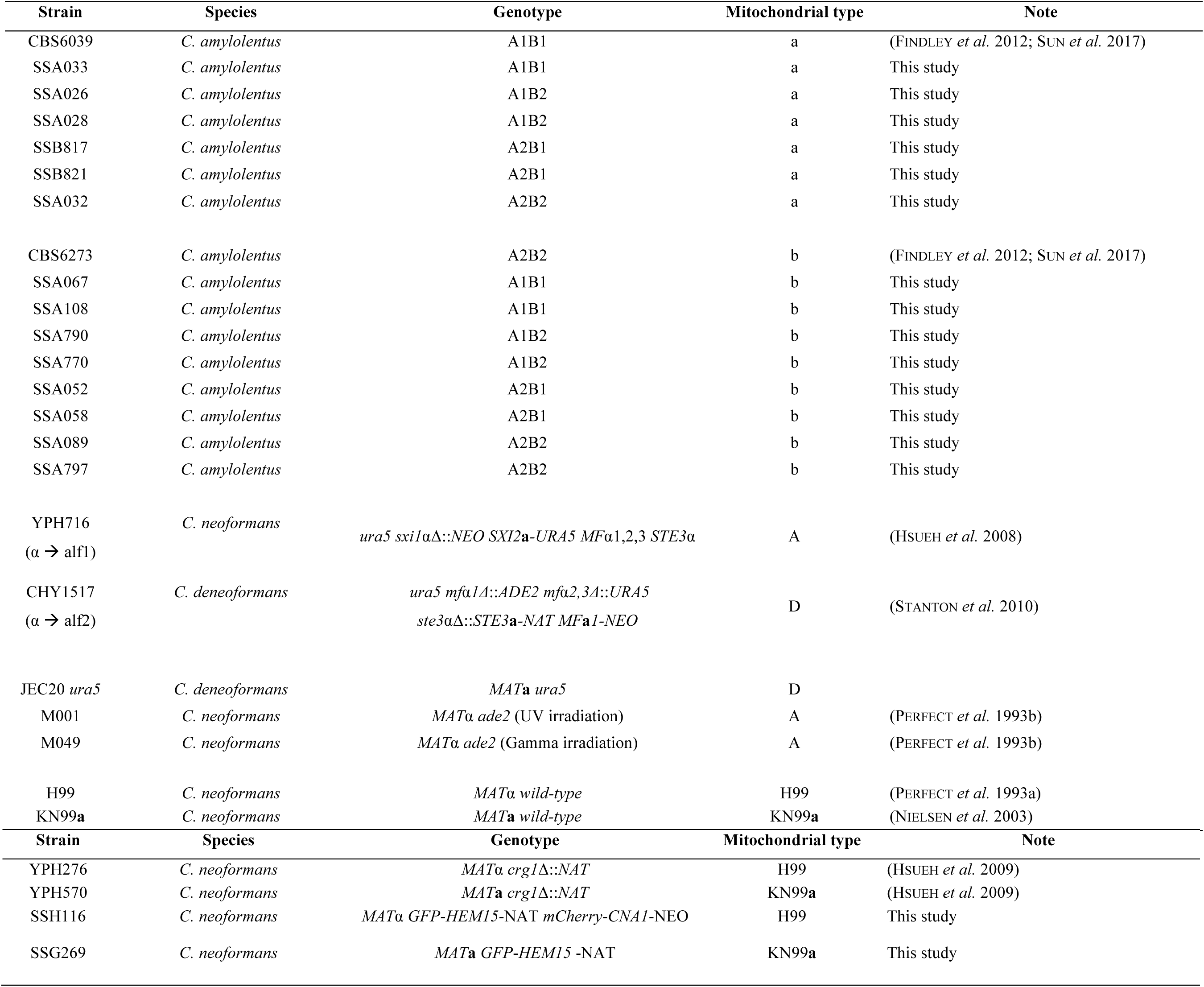
Strains analyzed in this study.

### Construction of deletion strains for testing the effects of individual genes on mito-UPI

For the bilateral crosses labeled as “H-K” in the Supplemental Table S2, deletion strains were constructed in the H99 (*MAT*α) and KN99**a** (*MAT***a**) backgrounds. For unilateral crosses labeled as “H-A”, such as those for the *MYO2* gene, the deletion strain was constructed by first generating a heterozygous deletion strain in the AI187 background. The AI187 heterozygous deletion strain was then induced to sporulate and the *MAT***a** progeny that inherited the deletion allele were recovered from randomly dissected basidiospores as described in previous studies. Mito-UPI was subsequently assessed in crosses between the *MAT***a** progeny with the deletion mutation and the Bt63 wildtype strain. For the genes *RCV1* and *YPT7*, the deletion strains were constructed in the KN99α background, and they were subsequently crossed with Bt63 and H99 wildtype strains, respectively, to assess the mito-UPI in these unilateral crosses.

### Laboratory crosses for analyzing mito-UPI in *C. amylolentus* and *C. neoformans*

For the strains used for crosses in *C. amylolentus*, other than the two natural isolates, CBS6039 and CBS6273, all of the other strains are meiotic progeny (basidiospores) dissected from crosses between CBS6039 and CBS6273. Three pairs of meiotic progeny, SSA026 and SSA028, SSA032 and SSA033, and SSA052 and SSA058, are each derived from three different basidia, and thus three independent meiotic events. All of the other progeny employed for crosses are from different basidia, thus representing independent meiotic events. Because we did not have selectable markers to select for fusion products in *C. amylolentus*, it is not known when mito-UPI is established during sexual reproduction. Therefore, we chose to dissect multiple spore chains representing independent meiotic events from each cross to interpret the pattern of mitochondrial inheritance and the origin of the mitochondria in the meiotic progeny.

For *C. neoformans*, mitochondrial inheritance was analyzed in three crosses between serotypes A and D isolates. First, mitochondrial inheritance was tested in a cross between strains CHY1517 and YPH716. CHY1517 has a serotype D *MAT*α background. While the *SXI1*α gene is intact, the *MAT*α genes encoding pheromones (*MF*α1, 2, 3) and pheromone receptors (*STE3*α) have been deleted, and replaced with *MAT***a** copies (*MF***a** and *STE3***a**). In addition, strain CHY1517 also has two phenotypic markers: *ura5* and NAT^R^ (Table 1; Figure 1) (STANTON *et al.* 2010). Strain YPH716 has a serotype A *MAT*α background, where the *SXI1*α gene was deleted, and a copy of the *SXI2***a** gene from the *MAT***a** background was transgenically introduced into its genome at the *URA5* locus (Table 1; Figure 1) (HSUEH *et al.* 2008). For comparison, mitochondrial inheritance was analyzed in two typical opposite sex crosses: JEC20 *ura5* × M001 (H99 *ade2*) and JEC20 *ura5* × M049 (H99 *ade2*). In these two crosses, the mitochondrial inheritance is expected to be uniparental from JOHE50 (*MAT***a**), based on previous reports (YAN AND XU 2003; YAN *et al.* 2007a).

**Figure 1.**
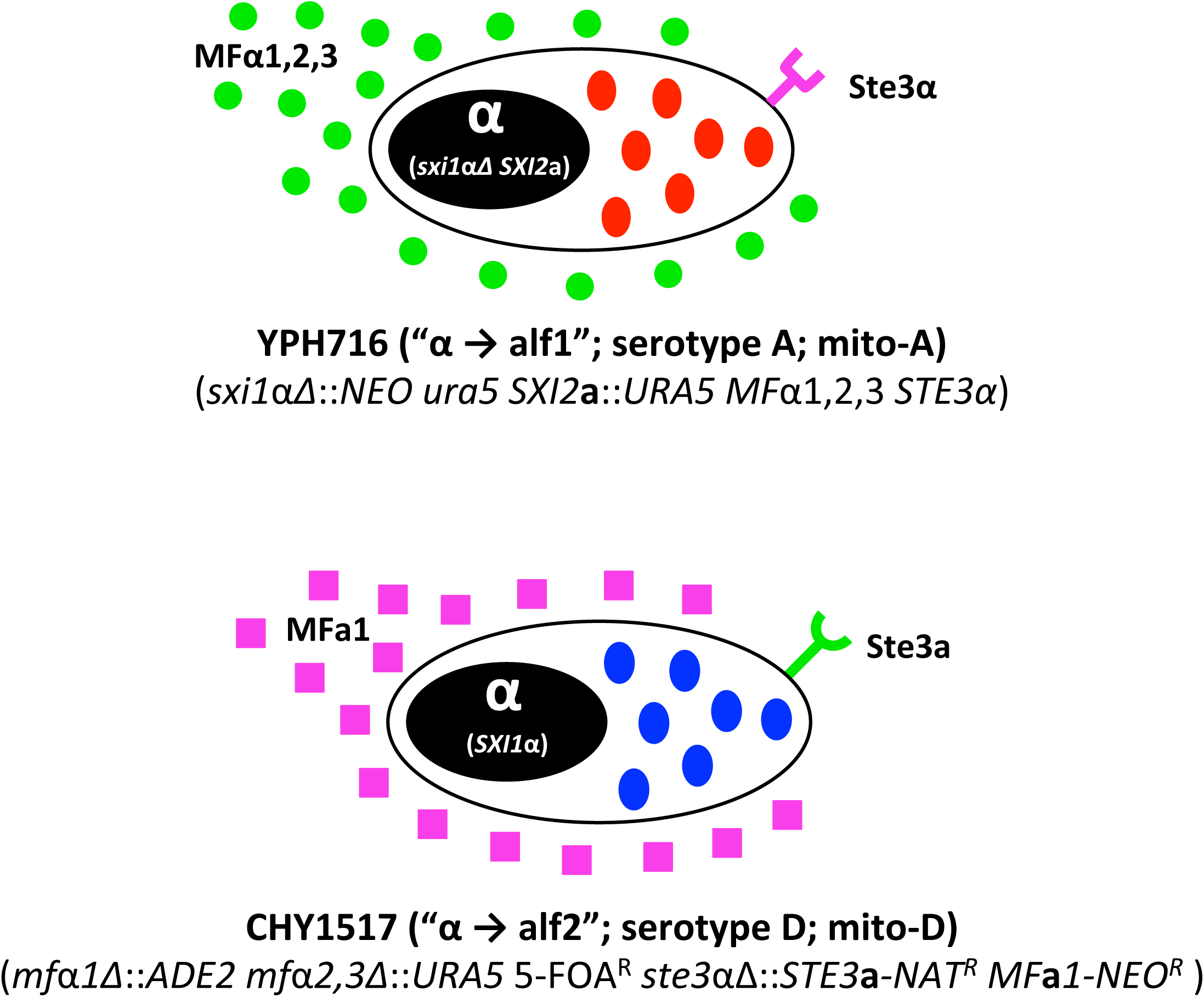
Illustration of the pheromones, pheromone receptors, and homeodomain transcription factors of the trans-mating type-engineered *Cryptococcus neoformans* strains. YPH716 (top) has a serotype A, *MAT*α genetic background, and serotype A mitochondria (red ovals). The *SXI1*α homeodomain transcription factor gene located in the *MAT* locus was deleted, and a copy of the *SXI2***a** gene was transgenically inserted at the *URA5* locus of the genome (HSUEH *et al.* 2008). Thus, during mating YPH716 expresses α pheromones and the *MF***a** pheromone receptor, but the **a** homeodomain transcription factor. CHY1517 (bottom) has a serotype D, *MAT*α genetic background, and serotype D mitochondria (blue ovals). The three α mating pheromone genes (*MF*α1, 2, and 3), as well as the α pheromone receptor (*STE*3α) were deleted from the *MAT* locus, and replaced with copies of the **a** mating pheromone (*MF***a**1) and pheromone receptor (*STE3***a**), respectively (STANTON *et al.* 2010). Thus, during mating CHY1517 expresses the **a** pheromone and *MF*α pheromone receptor, but the α homeodomain transcription factor. We designate YPH716 as “α **→** alf1” and CHY1517 as “α **→** alf2” to indicate the genetic modifications of their mating types, in which “alf1” and “alf2” designate the strains as “a-like faker” type 1 and type 2, respectively.

We also analyzed mitochondrial inheritance in unilateral and bilateral crosses of strains in which a single gene has been deleted (see Supplemental Table S2 for the list of genes). For these crosses, the parental strains are constructed in the H99 (*MAT*α) and KN99**a** (*MAT***a**) backgrounds, and thus, their mitochondrial types can be differentiated with PCR markers targeting the presence/absence of introns in the *COX1* gene as previously described (TOFFALETTI *et al.* 2004).

### Mating, spore dissection, and mating product screening

Mating and basidiospore dissection for *C. amylolentus* were carried out as previously described (FINDLEY *et al.* 2012). Briefly, mating compatible strains were mixed and spotted on V8 (pH=5) medium. The mating plates were incubated in the dark at room temperature (agar side up with no parafilm) for 1 – 2 weeks until abundant hyphae, basidia, and basidiospore chains were visible under the microscope. The basidiospore chains were then transferred onto fresh YPD medium, and individual basidiospores were separated using a fiber optic needle spore dissecting system, as previously described (IDNURM 2010). Individual spores separated from the same spore chain are the products of one meiotic event (FINDLEY *et al.* 2012).

For *C. neoformans*, the mating/fusion products from the cross between strains CHY1517 and YPH716 were obtained by first spotting the mixture of the two parental strains onto V8 (pH=5) medium, and then after hyphae, basidia, and basidiospore chains were formed, the hyphal sectors at the edge areas of the mating spots were excised and suspended in 1X PBS. The cell suspension was diluted, and spread onto SD-uracil plates to screen for Ura^+^ isolates. The Ura^+^ isolates were then transferred onto YPD+NAT plates to further screen for isolates that were also NAT resistant, and thus represented recombination/fusion of the markers present in the two parental strains.

Mating/fusion products from crosses between strains JEC20 *ura5* and M001 (H99 *ade2*) or JEC20 *ura5* and M049 (H99 *ade2*) were recovered similarly to those from the cross between CHY1517 and YPH716. Here, after the hyphal sectors were cut out and suspended in 1X PBS, the suspension was diluted and spread onto SD-uracil-adenine medium to screen for prototrophic isolates, and thus representing either recombinants or fusion products of the two parental strains.

To test the effect of a specific gene on mito-UPI, the unilateral and bilateral crosses were set up on MS media, incubated at room temperature in the dark for 10 days, and random spores were then dissected and analyzed as previously described (SUN *et al.* 2019).

### Genomic DNA, genetic markers, and genotyping

Germinated individual spores were transferred and patch-streaked onto fresh YPD medium, and genomic DNA was extracted from the biomass as described in a previous study (SUN *et al.* 2012). For *C. amylolentus*, mitochondrial genotyping was based on two PCR-RFLP markers targeting the *NAD4* and *NAD5* genes, respectively; while the mating types were determined using PCR-RFLP markers for *ETF1* and *SXI1* that are located within the A and B *MAT* locus, respectively (Supplemental Figure S1 and Supplemental Table S1). For *C. neoformans*, mitochondrial genotyping was based on the *NAD2* and *NAD5* genes, while serotype and mating-type specific markers for the *SXI1*α, *SXI2***a**, and *STE20*α/**a** genes were assayed to genotype the *MAT* locus (Supplemental Figure S1 and Supplemental Table S1). All of the markers are co-dominant (i.e. they can differentiate the two homozygous states, as well as the heterozygous one; Supplemental Figure S1). All PCR reactions were carried out using Promega 2X Go Taq Master Mix according to the manufacturer’s instructions, and with the following PCR thermal cycles: first an initial denaturation at 94°C for 6 minutes; then 36 cycles of 45 seconds at 94°C, 45 seconds at 60°C, and 90 seconds at 72°C; and a final extension at 72°C for 7 minutes. All enzyme digestions were performed with enzymes purchased from New England Biolabs and following the manufacturer’s instructions.

### Microscopy imaging

For fluorescence imaging of yeast cells, conjugation tubes, and initial hyphal formation, the mating mixture was grown on MS solid medium. The cells were incubated for 12 hours for yeast cells and conjugation tubes, and 24 hours for initial hyphal formation. The cells were collected, washed, and the cellular structures were observed and imaged with a ZEISS Imager widefield fluorescence microscope.

### Statistical analyses

Statistical tests of association of uniparental mitochondrial inheritance in *C. amylolentus* with specific mating-type locus/allele were carried out with a Binomial Probabilities Test (www.vassarstats.net). A P value of less than 0.05 is considered statistically significant, and is required to reject the null hypothesis that mitochondrial inheritance is not associated with a mating type locus/allele, i.e. the two mitochondrial types are inherited at equal frequency (50%) among all independent crosses, regardless of the mating types and mitochondrial types of the two parental strains. For mitochondrial inheritance in *C. neoformans*, Pearson’s chi-squared test (χ^2^ test) was applied to test whether the observed distribution among the progeny is significantly different from the null hypothesis of an equal chance of inheritance of the two mitochondrial types. P values of less than 0.05 were considered statistically significant and used to reject the null hypothesis.

### Data Availability

Strains and plasmids are available upon request. The authors affirm that all data necessary for confirming the conclusions of the article are present within the article, figures, and tables.

## RESULTS

### Mito-UPI in *C. amylolentus* is controlled by the *P/R MAT* locus

In a previous study, we demonstrated that *C. amylolentus* has a tetrapolar mating system with the two mating type loci, *P/R* locus (A locus) and *HD* locus (B locus), located on different chromosomes (FINDLEY *et al.* 2012). Additionally, we found that among all of the basidiospores generated by mating between the two natural *C. amylolentus* isolates, CBS6039 (A1B1, mito-a) and CBS6273 (A2B2, mito-b), mitochondria were always inherited from CBS6273, the A2B2 parent (FINDLEY *et al.* 2012). In the studies presented here, basidiospores produced by a CBS6039 × CBS6273 cross were dissected, and progeny with all four different mating types that have the mito-b mitochondrial type were recovered. Interestingly, during our efforts in dissecting additional basidiospores from the same cross, we found that out of the 40 basidiospore chains dissected and analyzed, basidiospores from two different spore chains all inherited mitochondria from CBS6039, the A1B1 parent with mitochondrial type mito-a (basidia No. 8 and No. 9 in Table 2). Thus, mitochondrial inheritance in *C. amylolentus* is indeed uniparental from the A2B2 parent (CBS6273), but with a low level of leakage (∼5%) from the other parent (CBS6039; A1B1). This low level of leakage allowed us to recover meiotic progeny of all four different mating types that possess the mito-a mitochondrial type.

**Table 2.** Summary of crosses and mitochondrial inheritance in *C. amylolentus*.

| Cross Type <sup>1</sup> | Cross <sup>1</sup> | Basidium | No. of<br>spores<br>dissected | No. of<br>spores<br>germinated | Germination<br>rate (%) |
| --- | --- | --- | --- | --- | --- |
| A1B1 (mito-a) × **<u>A2B2 (mito-b)</u>** | CBS6039 × **<u>CBS6273</u>** <sup>2</sup> | 1 | 16 | 6 | 38 |
|  |  | 2 | 15 | 8 | 53 |
|  |  | 3 | 13 | 6 | 46 |
|  |  | 4 | 25 | 13 | 52 |
|  |  | 5 | 17 | 8 | 47 |
|  |  | 6 | 12 | 5 | 42 |
|  |  | 7 | 18 | 16 | 89 |
|  |  | 8 <sup>3</sup> | 30 | 21 | 70 |
|  |  | 9 <sup>3</sup> | 26 | 18 | 69 |
|  | CBS6039 × **<u>SSA089</u>** | 1 | 19 | 10 | 53 |
|  |  | 2 | 14 | 10 | 71 |
|  | SSA033 × **<u>CBS6273</u>** | 1 | 17 | 14 | 82 |
|  |  | 2 | 15 | 12 | 80 |
|  | SSA033 × **<u>SSA089</u>** | 1 | 13 | 9 | 69 |
|  |  | 2 | 16 | 13 | 81 |
| A1B2 (mito-a) × **<u>A2B1 (mito-b)</u>** | SSA026 × **<u>SSA052</u>** | 1 | 20 | 5 | 25 |
|  | SSA026 × **<u>SSA058</u>** | 1 | 12 | 9 | 75 |
|  |  | 2 | 25 | 14 | 56 |
|  |  | 3 | 29 | 21 | 72 |
|  | SSA028 × **<u>SSA052</u>** | 1 | 12 | 7 | 58 |
|  | SSA028 × **<u>SSA058</u>** | 1 | 17 | 13 | 76 |
|  |  | 2 | 15 | 13 | 87 |
|  |  | 3 | 13 | 10 | 77 |
| **<u>A2B1 (mito-a)</u>** × A1B2 (mito-b) | **<u>SSB817</u>** × SSA770 | 1 | 6 | 4 | 67 |
|  |  | 2 | 8 | 2 | 25 |
|  |  | 3 | 8 | 3 | 38 |
|  |  | 4 | 14 | 9 | 64 |
|  | **<u>SSB817</u>** × SSA790 | 1 | 29 | 17 | 59 |
|  | **<u>SSB821</u>** × SSA797 | 1 | 11 | 5 | 45 |
|  |  | 2 | 15 | 3 | 20 |
| **<u>A2B2 (mito-a)</u>** × A1B1 (mito-b) | **<u>SSA032</u>** × SSA067 | 1 | 12 | 1 | 8 |
|  |  | 2 | 9 | 3 | 33 |
|  | **<u>SSA032</u>** × SSA108 | 1 | 7 | 4 | 57 |
|  |  | 2 | 11 | 10 | 91 |
|  |  | 3 | 12 | 4 | 33 |

<sup>1</sup>: The parental type highlighted in bold font and underlined is the parental type from which the meiotic progeny inherited the mitochondria.
<sup>2</sup>: Only nine of the more than 20 basidia that were dissected for crosses between CBS6039 and CBS6273 are shown here.
<sup>3</sup>: Basidia No. 8 and No. 9 inherited mito-a from the A1B1 parental strain, CBS6039.

We next conducted pairwise matings between isolates with compatible mating types but different mitochondrial types, and dissected multiple basidiospore chains from most of these crosses (Table 2). In all of these crosses, basidiospores dissected from the same cross always inherited the same mitochondria, further supporting previous observations that mitochondrial inheritance in *C. amylolentus* is predominantly UPI. Additionally, the mitochondria in all of the basidiospores were inherited from the parental strain that possessed the A2 *MAT* allele, independent of the alleles present at the B locus (Table 2). The observed association between the A2 allele and mitochondrial inheritance is statistically significant (Binomial Probability Test, P<0.05), and thus provides evidence that mito-UPI in *C. amylolentus* is controlled by the A2 mating type locus.

What gene(s) in the A2 locus is responsible for the mito-UPI in *C. amylolentus* then? As mentioned earlier, the A mating type locus in *C. amylolentus* has undergone extensive expansion compared to typical pheromone/pheromone receptor mating type loci in basidiomycetes, and contains ∼20 genes (FINDLEY *et al.* 2012; SUN *et al.* 2017). It is thus difficult to narrow down and define the genes that could be playing key roles in mito-UPI. However, we hypothesized that the pheromones and pheromone receptors might be involved in this process for the following reasons. First, pheromone and pheromone receptors are defining genes, and in some cases the only genes, of the *P/R* locus in basidiomycetes. If mito-UPI by the *P/R* locus is conserved in basidiomycetes, it is logical to hypothesize that the pheromones and pheromone receptors might be involved in this process. Also, pheromones and pheromone receptors are involved in the early stages of the mating process before zygote formation, including mating partner recognition and mating initiation. This coincides with the time when mitochondria from the two mating partners are thought to be differentially tagged for later protection or degradation in the zygotes. Additionally, in some basidiomycete species the expression of pheromones and pheromone receptors is not symmetrical between the mating partners (MCCLELLAND *et al.* 2004), which could provide opportunities for asymmetrical sexual development between the opposite mating types (please see below), as well as for the mitochondria of different mating types to be differentially tagged for protection/degradation, which collectively contribute to mito-UPI.

### Pheromone and pheromone receptors affect mito-UPI in *C. neoformans*

Because approaches for genetic manipulation of *C. amylolentus* are rudimentary, we decided to further investigate the role of pheromones and pheromone receptors in mito-UPI in its sister species, *C. neoformans*, based on the following considerations. First, *C. neoformans* is the most closely related known species to *C. amylolentus*. Additionally, it has been previously shown that mitochondria are uniparentally inherited in *C. neoformans* (XU *et al.* 2000; YAN AND XU 2003), and several genes involved in mating have been shown to play important roles in ensuring proper mito-UPI during mating (YAN *et al.* 2004; YAN *et al.* 2007a; GYAWALI AND LIN 2013). Given their close relationship, the mechanisms of mito-UPI are likely to be conserved between these two species. Second, although *C. neoformans* has a bipolar mating system governed by one large *MAT* locus, it has been shown that this large contiguous *MAT* locus is the result of a fusion event between the ancestral A and B loci, at which point the A locus had already undergone a similar expansion as in *C. amylolentus*, with most of the genes that have been identified within the A locus in *C. amylolentus* also present in the *MAT* locus of *C. neoformans* (HSUEH *et al.* 2011; FINDLEY *et al.* 2012; SUN *et al.* 2017).

To investigate the possible effects of pheromone and pheromone receptor genes on mito-UPI, two engineered *MAT*α strains were analyzed. In one strain, YPH716 (serotype A, “α **→ <u>a</u>**-<u>l</u>ike <u>f</u>aker 1 (alf1)”; Table 1; Figure 1), the original *SXI1*α gene in the *MAT*α locus has been deleted, and the *SXI2***a** from the *MAT***a** locus has been integrated at the *URA5* locus. Thus, YPH716 lacks the homeodomain transcription factor Sxi1α, and expresses instead the **a**-specific homeodomain transcription factor Sxi2**a**, along with all other α specific genes (Figure 1). In the second strain, CHY1517 (serotype D, “α **→** alf2”; Table 1; Figure 1), the *MAT*α genes encoding the pheromones (*MF*α1, *MF*α2, and *MF*α3) and pheromone receptor (*STE3*) have been deleted and replaced with the alleles from the *MAT***a** locus. As the result, CHY1517 expresses the Ste3**a** pheromone receptor and MF**a** instead of Ste3α and MFα, and has all other α-specific genes (Figure 1). Thus, if the null hypothesis that pheromones and pheromone receptors are not involved in mito-UPI is correct, we should expect during the mating between CHY1517 and YPH716 that the two types of mitochondria (mito-A and mito-D; Table 1 and Supplemental Figure S1) would be inherited at equal frequencies among meiotic progeny or fusion products of the two strains, as shown in a previous study (YAN *et al.* 2007a), because functional *SXI1* and *SXI2* genes are still present in the zygote. Alternatively, if pheromones and pheromone receptors are involved in mito-UPI, we would expect deviations of mitochondrial inheritance from biparental during the cross between CHY1517 and YPH716.

We recovered a total of 123 isolates from the cross between CHY1517 and YPH716 that have a combination of the phenotypic markers of the two parental strains (Table 3). Genotyping of these isolates using the four serotype and mating type specific markers (*SXI1*α, *SXI2***a**, and *STE20*α/**a**; Table 3) revealed that they all possess all four alleles, indicating that they are either serotype AD diploid fusion products of the two parental strains or meiotic products that are diploid or disomic for the *MAT* locus. Additionally, of these 123 mating/fusion products, 80 (65%) inherited mitochondria from CHY1517 (mito-D), 37 (30%) inherited mitochondria from YPH716 (mito-A), while 6 (5%) appear to have recombinant mitochondria (Table 3). The distribution of mito-A and mito-D among these mating products is significantly different from the null hypothesis that both mitochondrial types have equal chances (50:50) of inheritance (χ^2^ test, P<0.0001). Importantly, the biased mito-inheritance is in favor of those from the strain CHY1517, which has the *MF***a** and *STE3***a** alleles, although the other mating partner, YPH716, possesses the *SXI2***a** gene. Thus, our results suggest that the pheromones and pheromone receptors are indeed involved in controlling mito-UPI in *C. neoformans*. Specifically, replacing the *MAT*α version of pheromones and pheromone receptor with those from *MAT***a** in an otherwise *MAT*α strain significantly increased the chances of its mitochondria being inherited during sexual reproduction.

**Table 3.**
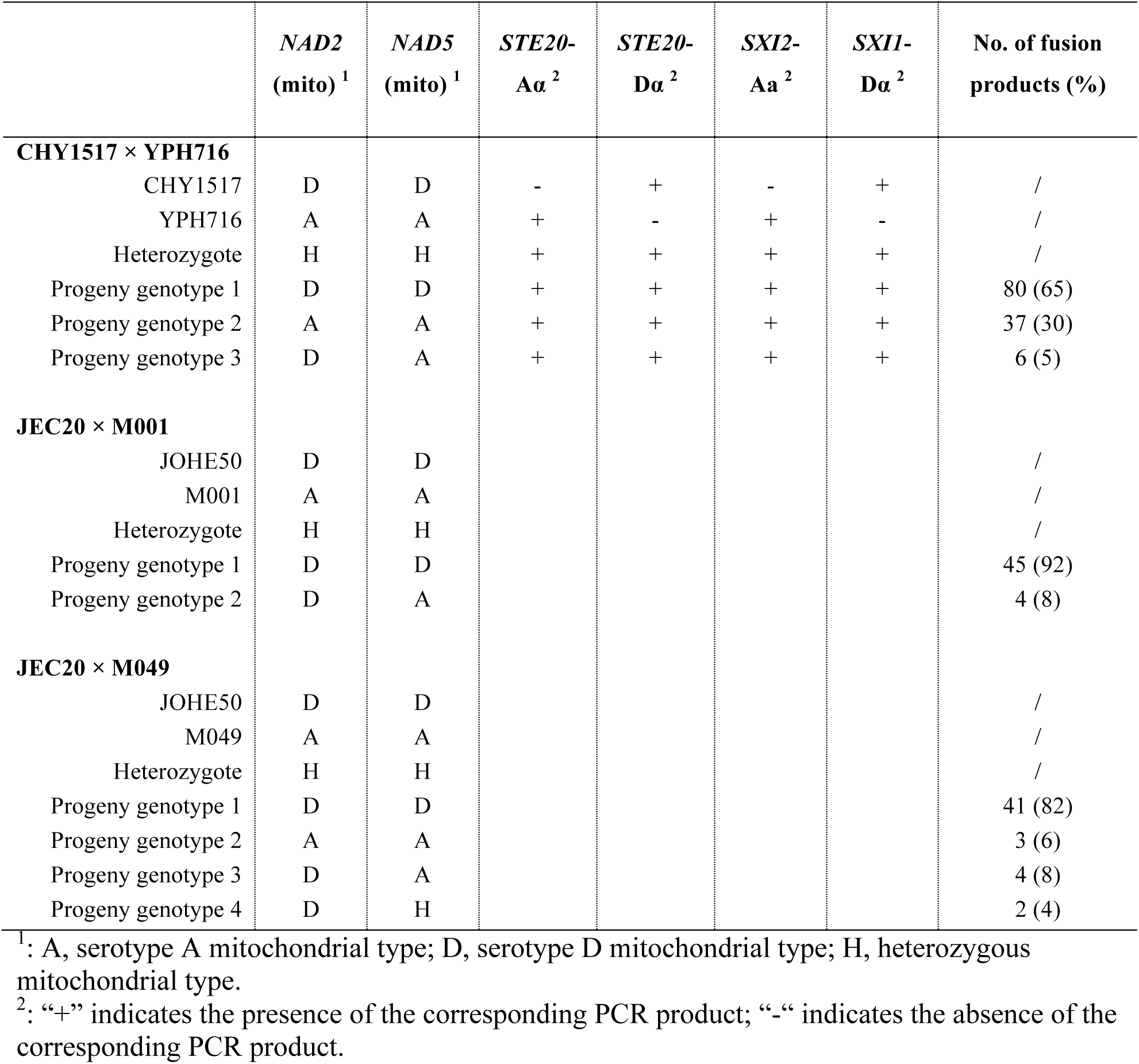
Summary of genotyping results of *C. neoformans* mating/fusion products.

However, it should be noted that mito-UPI is not fully restored to a wildtype level in the cross between CHY1517 and YPH716. For comparison, we performed two inter-serotype opposite-sex crosses using strains with intact mating type loci (JEC20 *ura5* × M001 (H99 *ade2*) and JEC20 *ura5* × M049 (H99 *ade2*); Table 3). In cross JEC20 *ura5* × M001, we found 45 out of 49 (92%) mating products inherited mitochondria from JEC20 *ura5*, the *MAT***a** parent, while the other 4 mating products (8%) inherited recombinant mitochondria. In cross JEC20 *ura5* × M049, we found that 41 out of 50 (82%) mating products inherited mitochondria from JEC20 *ura5*, 3 mating products (6%) inherited mitochondria from M049, and the other 6 mating products (12%) inherited recombinant mitochondria. In both cases, mito-UPI is occurring at a significantly higher frequency compared to the cross between CHY1517 and YPH716 (Fisher’s exact test, P<0.05). Taken together, these results suggest that pheromones and pheromone receptors contribute significantly to mito-UPI, but additional genes, which might also be located within the *MAT***a** locus, are also needed for wild-type levels of mito-UPI.

### Conjugation tube formation and hyphae initiation are asymmetrical in *C. neoformans* bisexual reproduction

In basidiomycetes, the interactions between pheromones and pheromone receptors are critical for the early stages of sexual reproduction, including mating partner sensing, conjugation formation, and zygote formation (BANDONI 1963; SPELLIG *et al.* 1994; BÖLKER 2001). To investigate the dynamics of sexual development and mitochondrial movement, we generated two *MAT***a** and *MAT*α strains, SSG269 and SSH116, respectively, in which the mitochondrial protein Hem15 was tagged with GFP in both strains. Additionally, for the *MAT*α strain (SSH116) the calcineurin A subunit Cna1 was tagged with mCherry. We set up crosses between these two strains and observed sexual development after 12 hours of mating on MS medium.

First, we found that the mitochondrial morphology is tubular when cells are growing on MS medium for 12 hours (Figure 2, I and II; >98%, n=50). This is consistent with previous studies showing tubulization of *C. neoformans* mitochondria under stress conditions (MA AND MAY 2010; CHANG AND DOERING 2018). Additionally, we observed that in cases where two cells are connected by a conjugation tube and the mCherry signal is not distributed universally throughout the “dumbbell” shaped structures, the conjugation tubes always have the mCherry signal (Figure 2, III; >98%, n=50). This suggests that during mating initiation, the conjugation tubes were always generated by the *MAT*α strain (SSH116) that is marked by the mCherry signal. There also appears to be separation between the mitochondria from the two parental cells and those in the conjugation tube (Figure 2, III; 80%, n = 15), suggesting that it is possible that during the initial stages of sexual development, the asymmetrical development of the conjugation tubes from the *MAT*α cells could generate a physical barrier that limits transmission of α mitochondria from the *MAT*α parent into the **a** cell side of the zygote.

**Figure 2.**
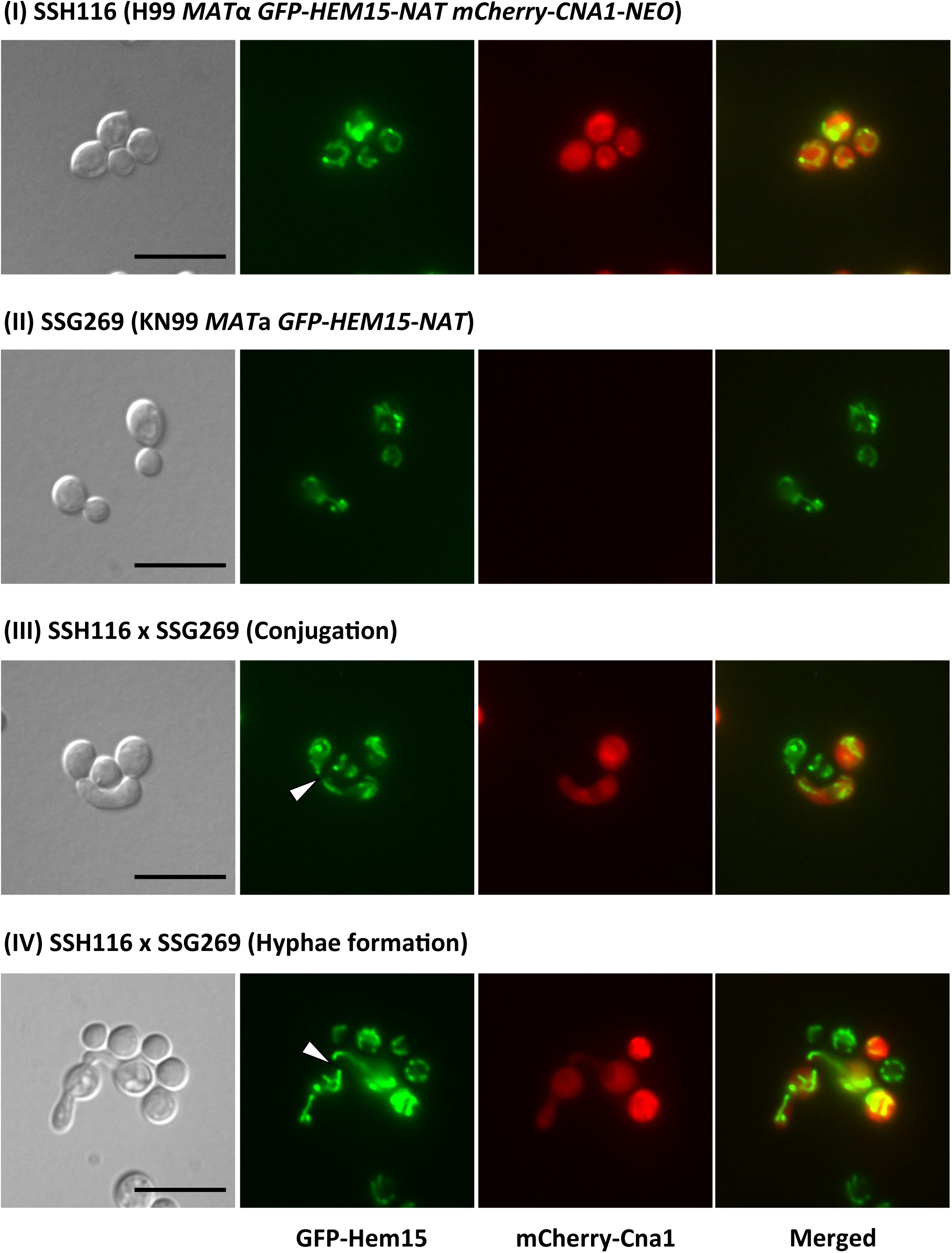
Microscope imaging of conjugation tube formation and hyphae initiation during *C. neoformans* sexual reproduction. For each panel, from left to right are DIC, GFP, mCherry, and merged images. (I) The microscopic images of the parental strain SSH116 after incubation of 12 hours on MS solid medium. (II) The microscopic images of the parental strain SSG269 after incubation of 12 hours on MS solid medium. (III) The microscopic images of the conjugation tube formation between strains SSH116 and SSG269 after 12 hours of coculturing on MS solid medium, in which the conjugation tubes are formed by the *MAT*α strain that has the mCherry signal. (IV) The microscopic images of the hyphae initiation in crosses between strains SSH116 and SSG269 after 24 hours of coculturing on MS solid medium, in which the hyphae are formed from the *MAT***a** cells, and on the opposite side of the conjugation tube. The white arrowheads highlight the gaps between mitochondrial complexes from the zygote and the conjugation tubes. Scale bar = 10 µm.

We also observed that after zygote formation (24 hours of mating on MS solid medium), the hyphae were almost always initiated from the zygote at a position opposite from the conjugation site, and the separation between the mitochondria from the zygote and the conjugation tube persisted (Figure 2, IV; 90%, n=10). This suggests that the mitochondria in the hyphae, and subsequently in the meiotic progeny, are mostly from the **a** cell side of the zygote, whose mitochondria are contributed mostly by the *MAT***a** parent.

### Deletion of the *CRG1* gene disrupts mito-UPI in bilateral crosses

It has been shown that the *CRG1* gene encodes a regulator of G-protein signaling (RGS) that negatively regulates the pheromone-signaling cascade and cAMP pathways during sexual reproduction in *C. neoformans*, with *crg1*Δ deletion strains displaying enhanced hyphal development in response to mating pheromones (FRASER *et al.* 2003; NIELSEN *et al.* 2003; WANG *et al.* 2004; FERETZAKI AND HEITMAN 2013). We hypothesized that deleting the *CRG1* gene might also disrupt the dynamics of pheromone and pheromone receptor interaction during early stages of mating when conjugation occurs, and consequently, compromise the fidelity of mito-UPI.

In confrontation assays involving the *crg1*Δ deletion strains, we found that both the *MAT*α and *MAT***a** deletion strains showed elevated pheromone response and filamentation compared to their respective wildtype strains (Figure 3A). However, this enhanced pheromone response is not symmetrical between the two mating types. Specifically, while the *MAT*α *crg1*Δ strain produced hyphae when confronted with the wildtype *MAT***a** as well as the *MAT***a** *crg1*Δ strains (Figure 3A, II and IV), the *MAT***a** *crg1*Δ strain only displayed enhanced pheromone response and hyphal growth when confronted with the *MAT*α *crg1*Δ strain but not the *MAT*α wildtype strain (Figure 3A, III and IV). Thus, it appears that the α mating type has an intrinsic higher sensitivity to pheromone compared to the **a** mating type.

**Figure 3.**
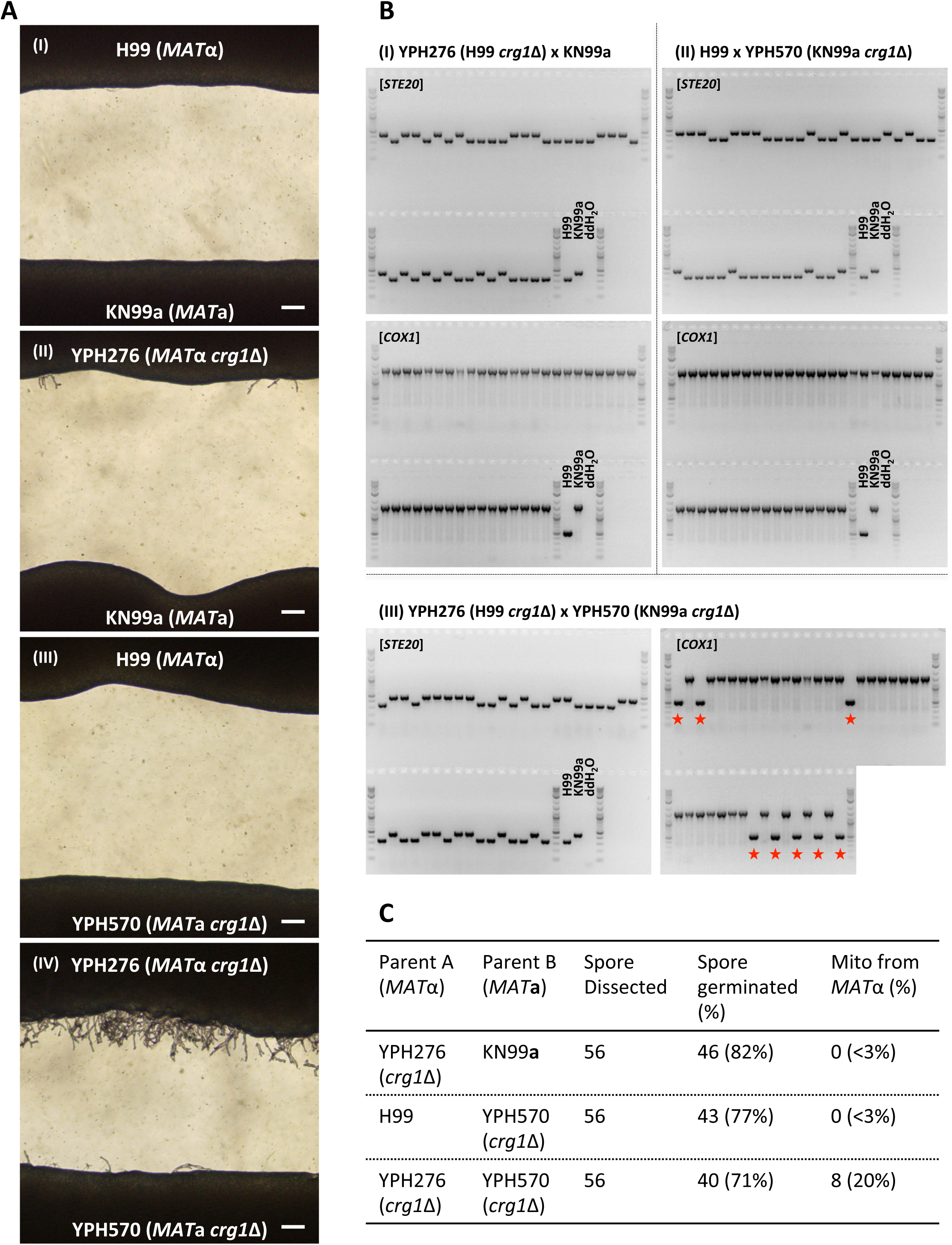
Effects of the *CRG1* gene on mating and mito-UPI. (A) Wildtype (I) as well as unilateral (II and III) and bilateral (IV) confrontation assays of *crg1*Δ deletion strains. Scale bar = 20 µm. (B) Genotyping of the mating type (*STE20*) and the mitochondrial type (*COX1*) of 40 random spores dissected from *crg1*Δ unilateral (I and II) and bilateral (III) crosses. The progeny highlighted with red stars are those that inherited mitochondria from the *MAT*α parent. (C) Summary of spore dissection, germination, and mito-UPI frequencies from *crg1*Δ unilateral and bilateral crosses.

We next dissected random basidiospores from the *crg1*Δ unilateral and bilateral crosses, and analyzed their mating types and mitochondrial types (Figure 3B). We found that while both unilateral crosses showed complete mito-UPI from the *MAT***a** parent in their progeny, progeny from the *crg1*Δ × *crg1*Δ bilateral cross showed a significantly higher level of mitochondrial leakage, with 8 of the 40 (20%) random basidiospores analyzed inheriting mitochondria from the *MAT*α parent (Figure 3C).

## DISCUSSION

Our data suggest that in addition to molecular mechanisms that actively degrade the mitochondria from the *MAT*α parent, the dynamics of early stages of sexual development controlled by the pheromone and pheromone receptors may also provide an additional physical barrier for the α mitochondrial to enter the zygote. Thus, consistent with previous hypotheses (e.g. in (WILSON AND XU 2012)), multiple mechanisms, both physically and genetically, could act in concert to ensure faithful mito-UPI from the *MAT***a** cell during *C. neoformans* sexual reproduction (Figure 4).

**Figure 4.**
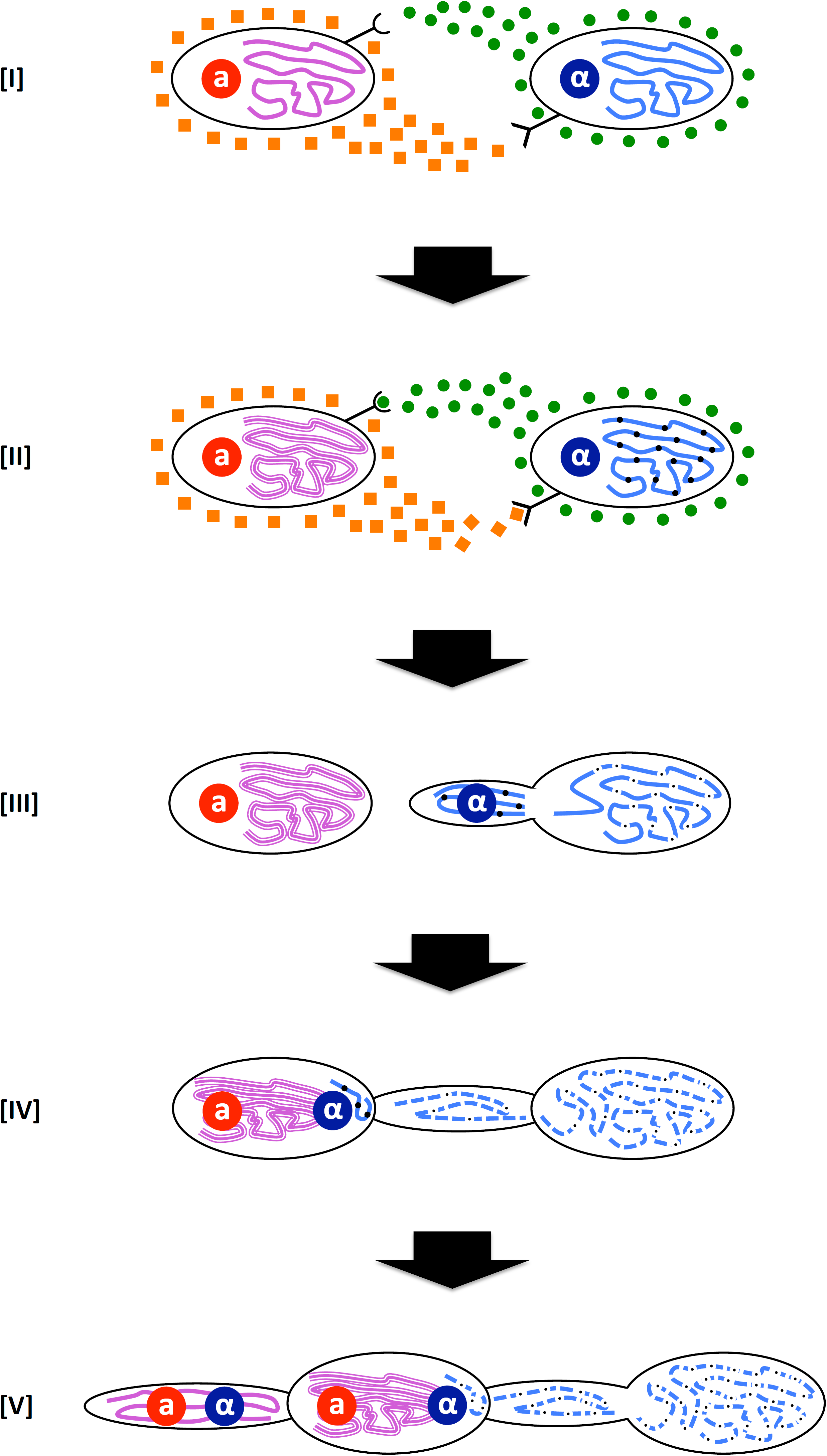
Model of mito-UPI regulation in *Cryptococcus* species. (I) and (II) The mito-UPI process starts when compatible mating partners sense pheromones from each other. The asymmetrical nature of this process could lead to activation of transcription factors (e.g. *MAT2*) and target genes whose products differentially tag the mitochondria in the two mating partners so that they would be targeted for or protected from degradation by mitophagy or other processes yet to be defined. (III) Successful pheromone and pheromone receptor interaction induces the formation of a conjugation tube from the *MAT*α parent, through which the *MAT*α nucleus migrates into the *MAT***a** parent to form the zygote. It is possible that the mitochondria remaining in the *MAT*α parent start degradation at this point, a process that could involve mitophagy. (IV) The zygote contains the two nuclei, as well as the mitochondria from the *MAT***a** parent. It is possible that trace amounts of mitochondria from the *MAT*α parent also migrate into the zygote. The mitochondria remaining in the conjugation tube and the *MAT*α parent cell continue degradation. (V) Hyphae are initiated from the zygote on the opposite side of the conjugation tube. This further increases the chances of mitochondria from the *MAT***a** parent being included in the hyphae, and consequently, the eventual mating progeny inherit mitochondria mostly from the *MAT***a** parent. Additionally, the trace amount of mitochondria from the *MAT*α parent would be degraded, through mechanisms such as mitophagy.

The mating process starts when compatible mating partners sense pheromones from each other. The asymmetrical nature of this process could lead to divergent activation of transcription factors (e.g. Mat2) and pathways (e.g. mitophagy), resulting in asymmetrical expression of genes critical for the initiation of sexual development (e.g. *STE3* and *MF*s) and differential tagging of the mitochondria in the two mating partners. Successful pheromone and pheromone receptor interaction induces the formation of conjugation tubes from the *MAT*α parent, through which the *MAT*α nucleus migrates into the *MAT***a** cell to form the zygote, while the vast majority of the α mitochondria remain in the *MAT*α cell and the conjugation tube, with only a small fraction migrating into the zygote accidentally. It is possible that the mitochondria remaining in the *MAT*α cell start degradation at this point, a process that could involve mitophagy, which could also direct the active degradation of mitochondria from the *MAT*α parent in the zygote. Subsequently, hyphae are initiated from the zygote on the opposite side of the conjugation tube, which could further limit the opportunities of mitochondria from the *MAT*α parent being included in the hyphae, even in cases where small amounts remain in the zygote at this stage. Consequently, the eventual mating progeny inherit mitochondria mostly from the *MAT***a** parent.

Our proposed model is consistent with current understandings of mito-UPI in *C. neoformans*. Studies of zygotes formed at the beginning of the mating process suggest that mito-UPI in *Cryptococcus* is likely established at an early stage of sexual development (SUN AND XU 2007; GYAWALI AND LIN 2013). In our studies of *C. amylolentus*, all of the basidiospores from the same basidium have identical mitochondrial type, and this is the case for all of the basidia, even the two anomalous basidia (#8 and #9) from the CBS6039 x CBS6273 crosses that exhibited mito-UPI exclusively from the A1 parent. In most genetic studies of fungal UPI in basidiomycetes, this has involved random spore dissection rather than dissection from individual basidia. For example the low level of UPI observed in *C. neoformans* from the α parent (generally <5%) has been termed “leakage” and has been thought to result from a low level of α mitochondria that survive in the hyphae. In this view, one might have expected to find ∼5% of spores dissected from an individual basidium would have the mitochondrial genome from the less favored parent. Instead, our observation suggests that mito-UPI may have been accomplished very early following zygote formation, such that ∼95% of the time the entire hyphal compartment has mitochondria derived from the A2 or the **a** parent. The remaining ∼<5% of the time the hyphal compartment would have the mitochondria derived from the A1 or the α parent. If the cost of inheriting more than one mitochondrial genotype is exerted on cell types beyond the zygote, such as in the hyphae, this may be a reason that the control of mito-UPI would be exerted early during sexual reproduction.

It has been shown that pheromones are asymmetrically expressed in the two mating types in *C. neoformans* under mating conditions, possibly through the regulation of Mat2, a key regulator of cell-cell fusion during zygote formation (SHEN *et al.* 2002; MCCLELLAND *et al.* 2004; KENT *et al.* 2008; LIN *et al.* 2010; GYAWALI AND LIN 2013). Specifically, the expression of mating pheromones from the *MAT***a** cells are increased earlier and to a higher extent than those from the *MAT*α cells (MCCLELLAND *et al.* 2004), suggesting that the pheromone sensing by the pheromone receptors, as well as the initiation of the downstream genes (e.g. mitophagy related mito-tagging) might also be asymmetrical between the two mating partners. We hypothesize that the observed asymmetrical gene expression underlies the observed unidirectional conjugation tube formation, as well as the polarized migration of the *MAT*α nucleus through the conjugation tube into the zygote, and thus these patterns of gene expression could be integral to mito-UPI. This is also consistent with our findings on the *crg1*Δ strains, where the deletion in the *MAT*α background exhibited elevated pheromone sensing and filamentation in a unilateral assay, while the *MAT***a** *crg1*Δ strain only showed enhanced pheromone response when confronted with the *MAT*α *crg1*Δ strain. This could be the reason why we failed to observe elevated mitochondrial leakage in the *crg1*Δ unilateral crosses, as the pheromone production and sensing dynamics in these cases are largely in accord with those present in the wildtype crosses. On the other hand, in *crg1*Δ bilateral crosses, because the *MAT***a** *crg1*Δ strain also showed elevated pheromone response and filamentation, the dynamics of pheromone production and response were thus disrupted and possibly as a consequence, increased levels of mitochondrial leakage in the basidiospores were observed. It should be noted that deletion of the *CRG1* gene appeared to have varied effects on mito-UPI in crosses between different species in the *C. gattii* species complex (WANG *et al.* 2015), suggesting the regulation of mito-UPI involves multiple genes and the process could be compromised during inter-species mating. This is also consistent with our analyses of the crosses between strains CHY1517 and YPH716. While CHY1517 possesses the *STE3***a** and *MF***a** genes, its mitochondria are not inherited at a level of 100% in the fusion products, suggesting that although the pheromone and pheromone receptor genes play important roles in controlling mito-UPI, other genes are required for faithful mito-UPI during sexual reproduction.

Functional homeodomain transcription factors *SXI1*α and *SXI2***a** from the *MAT*α and *MAT***a** parents, respectively, are both required for faithful mito-UPI in *C. neoformans* (YAN *et al.* 2004; YAN *et al.* 2007a), and it has been shown that deletion of the *SXI1*α gene enhances the spread of mobile mitochondrial introns in *C. neoformans* (YAN *et al.* 2018). The *SXI1*α*/SXI2***a** heterodimer serves as a key regulator of sexual development after zygote formation, including hyphal growth. Thus, it may play a critical role in determining the location of hyphal initiation to ensure it occurs away from the conjugation site, where occasional inclusion of *MAT*α mitochondria to the zygote might occur. For *C. amylolentus*, we showed that mito-UPI is controlled by the *P/R* locus. However, it should be noted that in this case the *HD* locus might still play a similar role as in *C. neoformans*. That is, functional *HD* heterodimers may be required to ensure the completion of mito-UPI. Because the strains that we used in *C. amylolentus* crosses are all derived from wild-type strains and possess functional *HD* genes, it would be interesting to test *C. amylolentus* crosses between isolates with mutated *HD* genes to see if faithful mito-UPI is still occurring.

Studies have shown that mitophagy is involved in uniparental mitochondrial inheritance in a variety of species (AL RAWI *et al.* 2011; LEVINE AND ELAZAR 2011; SATO AND SATO 2011; LUO *et al.* 2013), and it is hypothesized that differential tagging of mitochondria from the *MAT***a** and *MAT*α parents, and subsequent selective degradation of one group of mitochondria by mitophagy is also playing an important a role in mito-UPI in *C. neoformans* (GYAWALI AND LIN 2013). To investigate this, we deleted several known genes in the *S. cerevisiae* mitophagy pathway, as well as several other genes that are located within the *C. neoformans MAT* locus or have been shown to be involved in mitochondrial movement and segregation, and studied their effects on mito-UPI in both uni- and bi-lateral crosses (Supplemental Table S2). We did not find that deletion of any of these genes resulted in deviation from mito-UPI, suggesting these genes are not likely involved in mito-UPI. However, it is possible that other genes in these pathways that have not been tested are playing the key roles in mito-UPI, or that these genes are redundant with others not as yet tested. It is also possible that the keys genes involved in mito-UPI are pleiotropic and essential for cell survival, which would prevent them from being identified through gene deletion approaches.

Certain environmental factors, such as UV and temperature, influence mito-UPI in *C. neoformans* (YAN *et al.* 2007b), and natural strains of *C. gattii* with recombinant mitochondrial genomes have been isolated from the environment or following genetic crosses (VOELZ *et al.* 2013), suggestion mitochondrial leakage occurs during sexual reproduction in nature. It is possible that the expression of key genes involved in mito-UPI, such as the pheromone and pheromone receptor genes, as well as transcription factors including Mat2, Sxi1, and Sxi2, could be compromised by factors present in the natural environment, or become incompatible between mating partners due to sequence divergence accumulated between different lineages during evolution. Such environmental perturbance or gene function incompatibility could compromise both nuclear and mitochondrial dynamics during the early stages of sexual development, as well as the regulation of mitophagy pathway that is involved in mito-UPI, which collectively could lead to increased mitochondrial leakage from the *MAT*α parent.

Based on the DNA sequence, previous studies have shown that the *C. amylolentus* A1 pheromone receptor is more similar to the *STE3***a** allele in *C. neoformans*, and the A2 pheromone receptor is more similar to the *STE3*α allele. It is then intriguing that in *C. amylolentus* the mito--UPI is from the A2 parent, while in *C. neoformans* it is from the *STE3***a** parent. It should be noted that both the A1 and A2 *STE3* alleles show significant divergence from the *STE3***a** and *STE3*α alleles, respectively. Additionally, the *MAT* configurations in *C. amylolentus* and *C. neoformans* are highly different, with the former having a tetrapolar mating system with unlinked *P/R* and *HD* loci located on different chromosomes and the latter having a bipolar mating system with a fused *MAT* locus that contains both *P/R* and *HD* genes, demonstrating that the mating system has undergone significant transitions between the two species. It is possible that at some intermediate stages of this transition the mito-UPI was compromised. Because of the detrimental effects of mito-heteroplasy, selection pressure would have favored the re-establishment of mito-UPI. The linkage of this phenomenon to the mating type is the result of natural selection as this is the best way to ensure mito-UPI. During this re-establishment of mito-UPI, it is possible that the alleles associated with the mating type whose mitochondria are preferentially inherited in the progeny could change. Thus, while mito-UPI is the rule, its regulation could be dynamic, maybe particularly during speciation events.

## ACKNOWLEDGMENTS

We are grateful to Dr. Christina Hull for providing the genetically modified *C. neoformans* strain CHY1517 analyzed in this study. We thank Li Xu and Anna Floyd Averette for critical reading of the manuscript. This study was supported by NIH/NIAID R37 MERIT award AI39115-21 and R01 grant AI50113-15 awarded to J.H., and R01 grant AI133654-2 awarded to J.H., David Tobin, and Paul Magwene. J.H. is Co-Director and Fellow of CIFAR program Fungal Kingdom: Threats and Opportunities.

## SUPPLEMENTAL INFORMATION

**Figure S1.**
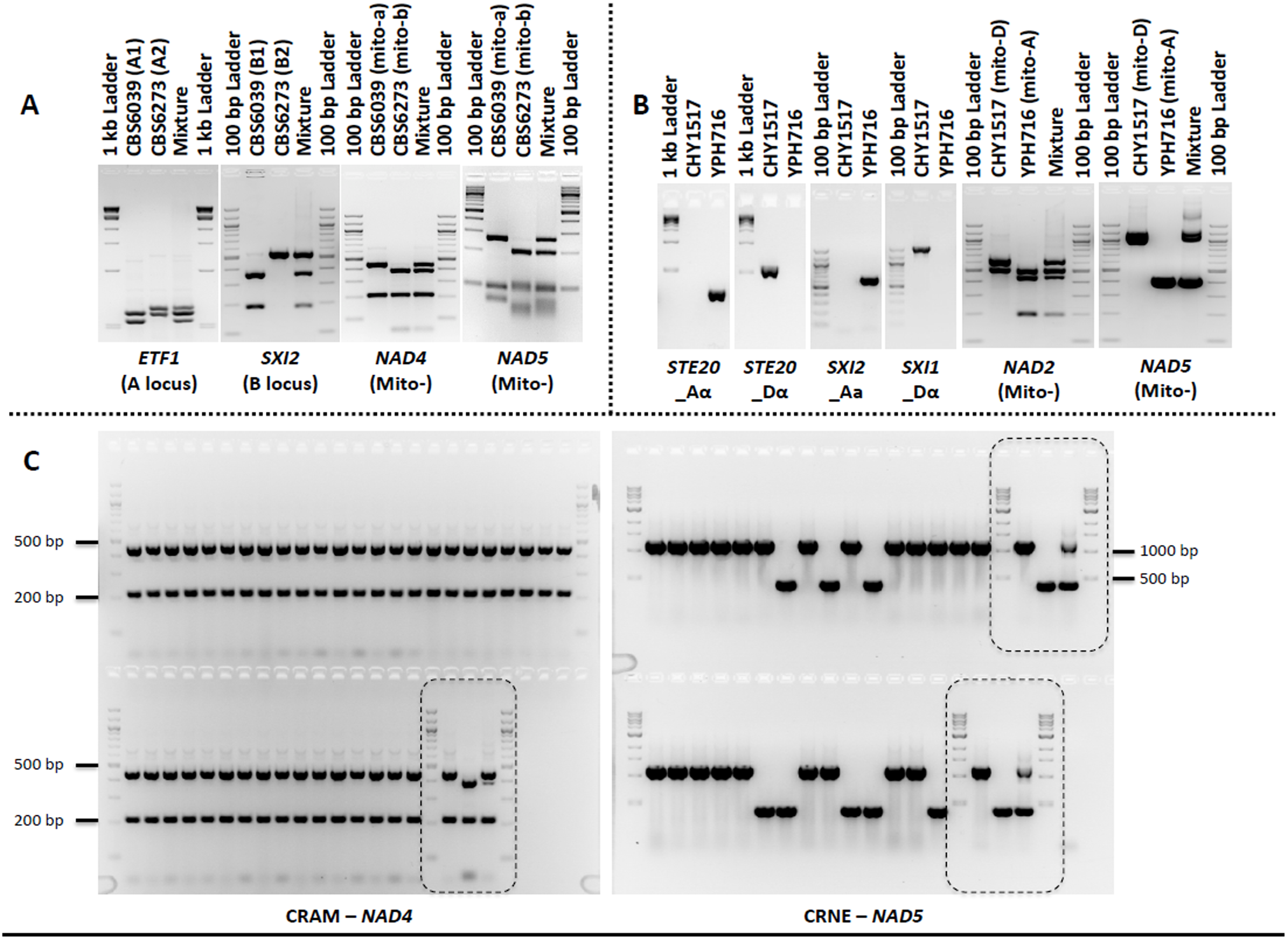
Nuclear and mitochondrial molecular markers used in this study. (A) Nuclear and mitochondrial markers employed for genotyping of *C. amylolentus*. (B) Nuclear and mitochondrial markers utilized for genotyping of *C. neoformans*. (C) Examples of mitochondrial genotyping of meiotic progeny from *C. amylolentus* (*NAD4*, left) and fusion products from *C. neoformans* (*NAD5*, right). The rectangles with dashed lines highlight the controls included during genotyping (in the same order as those shown in (A) and (B)). For all markers, “mixture” refers to an artificial heterozygote control by using a lab-generated mixture of two purified DNA samples that have different genotypes as the PCR template. Please see Supplemental Table S1 for detailed marker information. The unlabeled lanes in (A) and (B) are DNA marker ladders.

**Figure S2.**
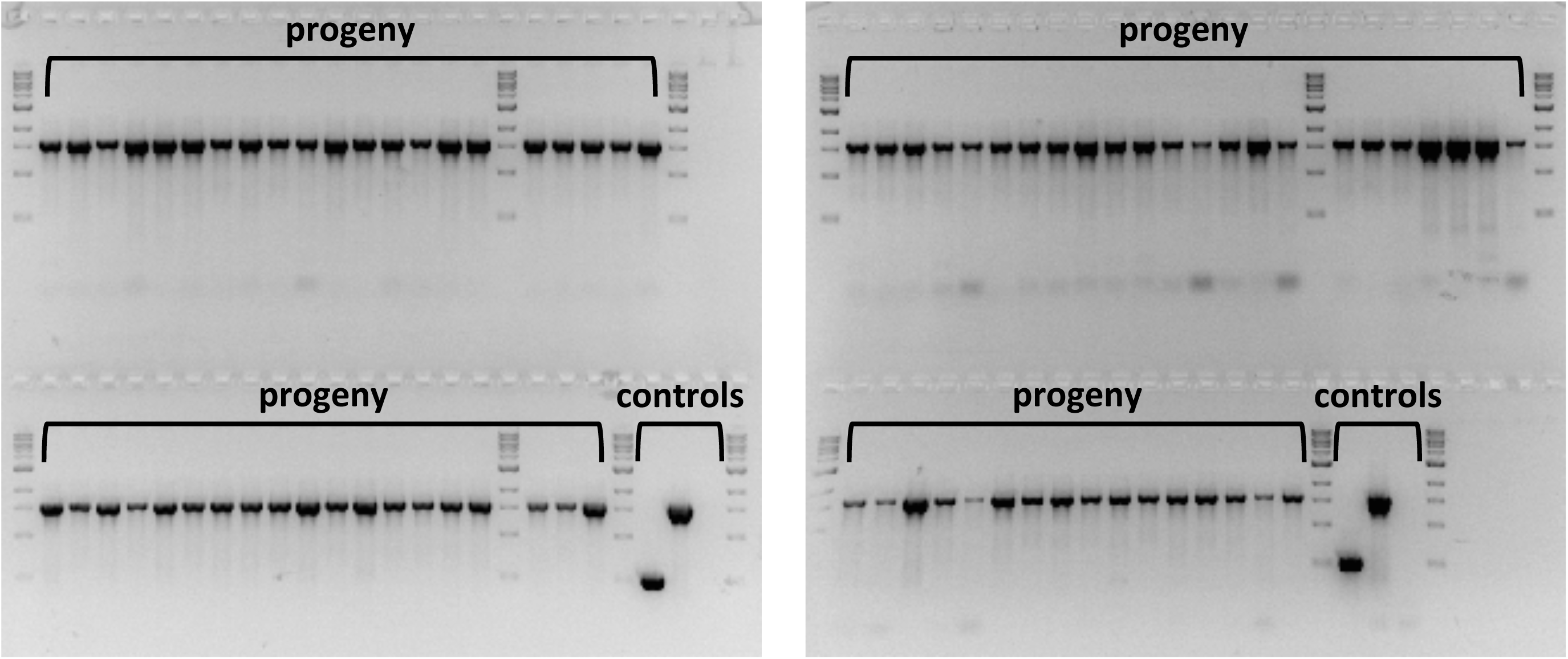
Progeny from *C. neoformans atg3*Δ and *atg7*Δ crosses showed mito-UPI. Mitochondrial genotyping of the *COX1* locus showed that progeny from the *atg3*Δ (left, 40 random basidiospores) and *atg7*Δ (right, 39 random spores) crosses exhibited mitochondrial UPI. Controls from left to right are: *MAT*α parent, *MAT***a** parent, and ddH_2_O, respectively.

**Table S1.**
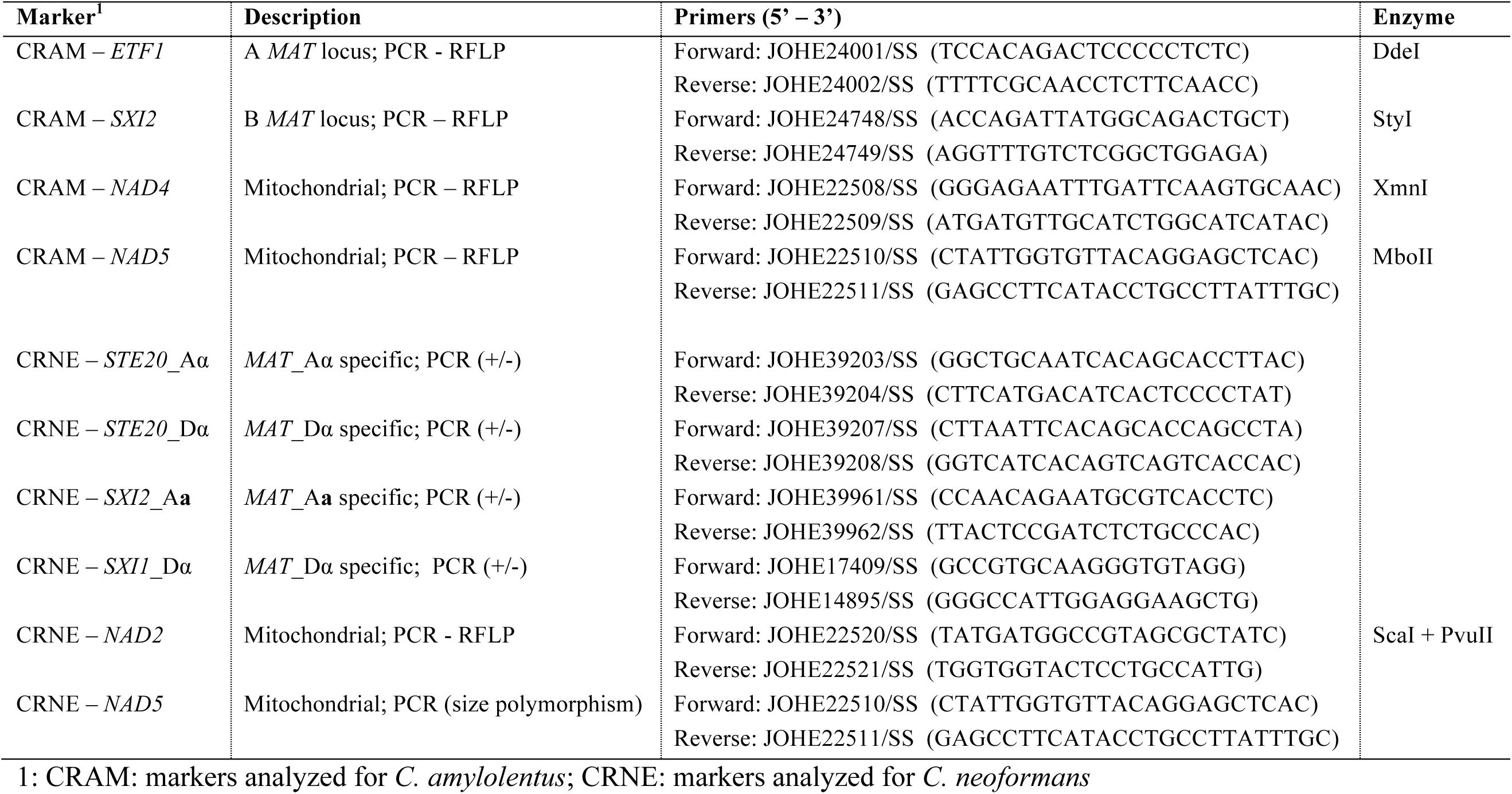
Markers and primers employed in this study.

**Table S2.** Genes tested for their effects on mito-UPI in unilateral and bilateral crosses.

| **Group** | **Gene** | **ID (in H99)** | **Strain (*MAT*)** | **Crosses** | **Backgrounds<sup>a</sup>** |
| --- | --- | --- | --- | --- | --- |
| Within the *MAT* locus | *SPO14* | CNAG_06812 | SSC927 ($\alpha$)<br>SSC937 (**a**) | Bilateral | H-K |
| | *ETF1* | CNAG_06806 | SSE720 ($\alpha$)<br>SSE693 (**a**) | Bilateral | H-K |
| | *RUM1* | CNAG_07411 | SSC919 ($\alpha$)<br>SSC938 (**a**) | Bilateral | H-K |
| | *CAP1* | CNAG_06813 | SSC929 ($\alpha$)<br>SSC940 (**a**) | Bilateral | H-K |
| | *BSP1* | CNAG_01457 | SSC939 ($\alpha$)<br>SSC941 (**a**) | Bilateral | H-K |
|  | *MYO2* | CNAG_06971 |  | Unilateral | H-A |
| Mitophagy related | *ATG1* | CNAG_05005 | SSH126 ($\alpha$)<br>SSH159 (**a**) | Bilateral | H-K |
| | *ATG3* | CNAG_06892 | SSG669 ($\alpha$)<br>SSH298 (**a**) | Bilateral | H-K |
| | *ATG7* | CNAG_04538 | SSG673 ($\alpha$)<br>SSH302 (**a**) | Bilateral | H-K |
| | *ATG12* | CNAG_07645 | SSH129 ($\alpha$)<br>SSH153 (**a**) | Bilateral | H-K |
| | *PTC6* | CNAG_06418 | SSG923 ($\alpha$)<br>SSG930 (**a**) | Bilateral | H-K |
| | *VPS1* | CNAG_00565 | SSG926 ($\alpha$)<br>SSG909 (**a**) | Bilateral | H-K |
| Involved in mitochondrial movement and segregation | *MMM1* | CNAG_05557 | Ianiri and Idnurm, 2013 | Unilateral | H-A |
|  | *MDM10* | CNAG_06345 | Ianiri and Idnurm, 2013 | Unilateral | H-A |
|  | *MDM34* | CNAG_02304 | Ianiri and Idnurm, 2013 | Unilateral | H-A |

| Group | Gene | ID (in H99) | Strain (*MAT*) | Crosses | Backgrounds <sup>a</sup> |
| --- | --- | --- | --- | --- | --- |
|  | *RCV1* | CNAG_01983 | Jung et al., 2015 | Unilateral | H-B |
|  | *YPT7* | CNAG_02575 | Jung et al., 2015 | Unilateral | H-K |

<sup>a</sup>: The effects of mito-UPI are tested in crosses between H99 (*MAT* $\alpha$ , H) with *MATa* strains KN99a (K), Bt63a (B), or the *MATa* progeny derived from the diploid strain AI187 that inherited the deletion allele (A). All analyses were based on a minimum of 40 randomly dissected basidiospores.

## REFERENCES

Al Rawi, S., S. Louvet-Vallée, A. Djeddi, M. Sachse, E. Culetto et al., 2011 Postfertilization autophagy of sperm organelles prevents paternal mitochondrial DNA transmission. Science 334: 1144–1147.

Bandoni, R. J., 1963 Conjugation in *Tremella mesenterica*. Canadian Journal of Botany 41: 467–474.

Barr, C. M., M. Neiman and D. R. Taylor, 2005 Inheritance and recombination of mitochondrial genomes in plants, fungi and animals. New Phytologist 168: 39–50.

Berger, K. H., and M. P. Yaffe, 2000 Mitochondrial DNA inheritance in *Saccharomyces cerevisiae*. Trends in Microbiology 8: 508–513.

Birky, C., 1996 Uniparental inheritance of mitochondrial and chloroplast genes: Mechanisms and evolution. Proc. Natl. Acad. Sci. USA 92: 11331.

Birky, C. W., 2001 The inheritance of genes in mitochondria and chloroplasts: laws, mechanisms, and models. Annual Review of Genetics 35: 125–148.

Bölker, M., 2001 *Ustilago maydis*-a valuable model system for the study of fungal dimorphism and virulence. Microbiology 147: 1395–1401.

Bortfeld, M., K. Auffarth, R. Kahmann and C. W. Basse, 2004 The *Ustilago maydis* a2 mating-type locus genes *lga2* and *rga2* compromise pathogenicity in the absence of the mitochondrial p32 family protein Mrb1. Plant Cell 16: 2233–2248.

Chang, A. L., and T. L. Doering, 2018 Maintenance of mitochondrial morphology in *Cryptococcus neoformans* is critical for stress resistance and virulence. mBio 9: e01375–01318.

Egal, R., J. Kohli, P. Thuriaux and K. Wolf, 1980 Genetics of the fission yeast *Schizosaccharomyces pombe*. Annual Review of Genetics 14: 77–108.

Fedler, M., K.-S. Luh, K. Stelter, F. Nieto-Jacobo and C. W. Basse, 2009 The *a2* mating-type locus genes *lga2* and *rga2* direct uniparental mitochondrial DNA (mtDNA) inheritance and constrain mtDNA recombination during sexual development of *Ustilago maydis*. Genetics 181: 847–860.

Feretzaki, M., and J. Heitman, 2013 Genetic circuits that govern bisexual and unisexual reproduction in *Cryptococcus neoformans*. PLoS Genetics 9: e1003688.

Findley, K., M. Rodriguez-Carres, B. Metin, J. Kroiss, A. Fonseca et al., 2009 Phylogeny and phenotypic characterization of pathogenic *Cryptococcus* species and closely related saprobic taxa in the *Tremellales*. Eukaryotic Cell 8: 353–361.

Findley, K., S. Sun, J. A. Fraser, Y.-P. Hsueh, A. F. Averette et al., 2012 Discovery of a modified tetrapolar sexual cycle in *Cryptococcus amylolentus* and the evolution of *MAT* in the *Cryptococcus* species complex. PLoS Genetics 8: e1002528.

Fraser, J. A., R. L. Subaran, C. B. Nichols and J. Heitman, 2003 Recapitulation of the sexual cycle of the primary fungal pathogen *Cryptococcus neoformans* var. *gattii*: implications for an outbreak on Vancouver island, Canada. Eukaryotic Cell 2: 1036–1045.

Gardner, A., and R. G. Boles, 2005 Is a “Mitochondrial Psychiatry” in the future? A review. Current Psychiatry Reviews 1: 255–271.

Goodenough, U., and J. Heitman, 2014 Origins of eukaryotic sexual reproduction. Cold Spring Harbor Perspectives in Biology 6: a016154.

Gyawali, R., and X. Lin, 2011 Mechanisms of uniparental mitochondrial DNA inheritance in *Cryptococcus neoformans*. Mycobiology 39: 235–242.

Gyawali, R., and X. Lin, 2013 Prezygotic and postzygotic control of uniparental mitochondrial DNA inheritance in *Cryptococcus neoformans*. mBio 4: e00112–00113.

Hagen, F., K. Khayhan, B. Theelen, A. Kolecka, I. Polacheck et al., 2015 Recognition of seven species in the *Cryptococcus gattii*/*Cryptococcus neoformans* species complex. Fungal Genetics and Biology 78: 16–48.

Hintz, W., J. B. Anderson and P. A. Horgen, 1988 Nuclear migration and mitochondrial inheritance in the mushroom *Agaricus bitorquis*. Genetics 119: 35–41.

Hsueh, Y. P., J. A. Fraser and J. Heitman, 2008 Transitions in sexuality: recapitulation of an ancestral tri- and tetrapolar mating system in *Cryptococcus neoformans*. Eukaryotic Cell 7: 1847–1855.

Hsueh, Y. P., B. Metin, K. Findley, M. Rodriguez-Carres and J. Heitman, 2011 The mating type locus of *Cryptococcus*: evolution of gene clusters governing sex determination and sexual reproduction from the phylogenomic perspective, pp. 139-149 in Cryptococcus: from human pathogen to model yeast, edited by J. Heitman, T. R. Kozel, K. J. Kwon-Chung, J. R. Perfect and A. Casadevall. American Society for Microbiology, Washington, DC.

Hsueh, Y. P., C. Xue and J. Heitman, 2009 A constitutively active GPCR governs morphogenic transitions in *Cryptococcus neoformans*. EMBO J 28: 1220–1233.

Idnurm, A., 2010 A tetrad analysis of the basidiomycete fungus *Cryptococcus neoformans*. Genetics 185: 153–163.

Ingavale, S. S., Y. C. Chang, H. Lee, C. M. McClelland, M. L. Leong et al., 2008 Importance of mitochondria in survival of *Cryptococcus neoformans* under low oxygen conditions and tolerance to cobalt chloride. PLoS Pathog 4: e1000155.

Kent, C. R., P. Ortiz-Bermúdez, S. S. Giles and C. M. Hull, 2008 Formulation of a defined V8 medium for induction of sexual development of *Cryptococcus neoformans*. Applied and Environmental Microbiology 74: 6248–6253.

Kretschmer, M., J. Wang and J. W. Kronstad, 2012 Peroxisomal and mitochondrial β-oxidation pathways influence the virulence of the pathogenic fungus *Cryptococcus neoformans*. Eukaryotic Cell 11: 1042–1054.

Kwon-Chung, K. J., 1975 A new genus, *Filobasidiella*, the perfect state of *Cryptococcus neoformans*. Mycologia 67: 1197–1200.

Lane, N., 2012 The problem with mixing mitochondria. Cell 151: 246-248.

Lesnefsky, E. J., S. Moghaddas, B. Tandler, J. Kerner and C. L. Hoppel, 2001 Mitochondrial dysfunction in cardiac disease: ischemia -- reperfusion, aging, and heart failure. Journal of Molecular and Cellular Cardiology 33: 1065–1089.

Levine, B., and Z. Elazar, 2011 Inheriting maternal mtDNA. Science 334: 1069–1070.

Lin, X., J. C. Jackson, M. Feretzaki, C. Xue and J. Heitman, 2010 Transcription factors Mat2 and Znf2 operate cellular circuits orchestrating opposite- and same-sex mating in *Cryptococcus neoformans*. PLOS Genetics 6: e1000953.

Loftus, B. J., E. Fung, P. Roncaglia, D. Rowley, P. Amedeo et al., 2005 The genome of the basidiomycetous yeast and human pathogen *Cryptococcus neoformans*. Science 307: 1321–1324.

Luo, S.-M., Z.-J. Ge, Z.-W. Wang, Z.-Z. Jiang, Z.-B. Wang et al., 2013 Unique insights into maternal mitochondrial inheritance in mice. Proceedings of the National Academy of Sciences, USA 110: 13038–13043.

Ma, H., F. Hagen, D. J. Stekel, S. A. Johnston, E. Sionov et al., 2009 The fatal fungal outbreak on Vancouver Island is characterized by enhanced intracellular parasitism driven by mitochondrial regulation. Proc Natl Acad Sci U S A 106: 12980–12985.

Ma, H., and R. C. May, 2010 Mitochondria and the regulation of hypervirulence in the fatal fungal outbreak on Vancouver Island. Virulence 1: 197–201.

Maheshwari, S., and D. A. Barbash, 2011 The genetics of hybrid incompatibilities. Annual Review of Genetics 45: 331–355.

Mahlert, M., C. Vogler, K. Stelter, G. Hause and C. W. Basse, 2009 The *a2* mating-type-locus gene *lga2* of *Ustilago maydis* interferes with mitochondrial dynamics and fusion, partially in dependence on a Dnm1-like fission component. Journal of Cell Science 122: 2402–2412.

May, G., and J. W. Taylor, 1988 Patterns of mating and mitochondrial DNA inheritance in the agaric basidiomycete *Coprinus cinereus*. Genetics 118: 213–220.

McBride, H. M., M. Neuspiel and S. Wasiak, 2006 Mitochondria: more than just a powerhouse. Current Biology 16: R551–R560.

McClelland, C. M., Y. C. Chang, A. Varma and K. J. Kwon-Chung, 2004 Uniqueness of the mating system in *Cryptococcus neoformans*. Trends Microbiol. 12: 208–212.

Ni, M., M. Feretzaki, S. Sun, X. Wang and J. Heitman, 2011 Sex in Fungi. Annual Review of Genetics 45: 405–430.

Nielsen, K., G. M. Cox, P. Wang, D. L. Toffaletti, J. R. Perfect et al., 2003 Sexual cycle of *Cryptococcus neoformans* var. *grubii* and virulence of congenic a and α isolates. Infect Immun 71: 4831–4841.

Passer, A. R., M. A. Coelho, R. B. Billmyre, M. Nowrousian, M. Mittelbach et al., 2019 Genetic and genomic analyses reveal boundaries between species closely related to *Cryptococcus* pathogens. mBio 10: e00764–00719.

Perfect, J. R., N. Ketabchi, G. M. Cox, C. W. Ingram and C. L. Beiser, 1993a Karyotyping of *Cryptococcus neoformans* as an epidemiological tool. Journal of Clinical Microbiology 31: 3305–3309.

Perfect, J. R., D. L. Toffaletti and T. H. Rude, 1993b The gene encoding phosphoribosylaminoimidazole carboxylase (*ADE2*) is essential for growth of *Cryptococcus neoformans* in cerebrospinal fluid. Infect. Immun. 61: 4446–4451.

Sato, M., and K. Sato, 2011 Degradation of paternal mitochondria by fertilizationtriggered autophagy in *C. elegans* embryos. Science 334: 1141–1144.

Shakya, V. P. S., and A. Idnurm, 2013 Sex determination directs uniparental mitochondrial inheritance in *Phycomyces*. Eukaryotic Cell: EC.00203-00213.

Sharpley, M. S., C. Marciniak, K. Eckel-Mahan, M. McManus, M. Crimi et al., 2012 Heteroplasmy of mouse mtDNA is genetically unstable and results in altered behavior and cognition. Cell 151: 333–343.

Shen, W. C., R. C. Davidson, G. M. Cox and J. Heitman, 2002 Pheromones stimulate mating and differentiation via paracrine and autocrine signaling in *Cryptococcus neoformans*. Eukaryot. Cell 1: 366–377.

Shingu-Vazquez, M., and A. Traven, 2011 Mitochondria and fungal pathogenesis: drug tolerance, virulence, and potential for antifungal therapy. Eukaryotic Cell 10: 1376–1383.

Spellig, T., M. Bolker, F. Lottspeich, R. W. Frank and R. Kahmann, 1994 Pheromones trigger filamentous growth in *Ustilago maydis*. Embo J. 13: 1620–1627.

Stanton, B. C., S. S. Giles, M. W. Staudt, E. K. Kruzel and C. M. Hull, 2010 Allelic exchange of pheromones and their receptors reprograms sexual identity in *Cryptococcus neoformans*. PLoS Genet 6: e1000860.

Sun, S., Y.-P. Hsueh and J. Heitman, 2012 Gene conversion occurs within the mating-type locus of *Cryptococcus neoformans* during sexual reproduction. PLoS Genetics 8: e1002810.

Sun, S., S. J. Priest and J. Heitman, 2019 *Cryptococcus neoformans* mating and genetic crosses. Current Protocols in Microbiology 53: e75.

Sun, S., and J. Xu, 2007 Genetic analyses of a hybrid cross between serotypes A and D strains of the human pathogenic fungus *Cryptococcus neoformans*. Genetics 177: 1475–1486.

Sun, S., V. Yadav, R. B. Billmyre, C. A. Cuomo, M. Nowrousian et al., 2017 Fungal genome and mating system transitions facilitated by chromosomal translocations involving intercentromeric recombination. PLoS Biology 15: e2002527.

Toffaletti, D. L., K. Nielsen, F. Dietrich, J. Heitman and J. R. Perfect, 2004 *Cryptococcus neoformans* mitochondrial genomes from serotype A and D strains do not influence virulence. Curr Genet 46: 193–204.

Voelz, K., H. Ma, S. Phadke, E. J. Byrnes, P. Zhu et al., 2013 Transmission of hypervirulence traits via sexual reproduction within and between lineages of the human fungal pathogen *Cryptococcus gattii*. PLOS Genetics 9: e1003771.

Wagner-Vogel, G., F. Lämmer, J. Kämper and C. W. Basse, 2015 Uniparental mitochondrial DNA inheritance is not affected in *Ustilago maydis Δatg11* mutants blocked in mitophagy. BMC Microbiology 15: 23.

Wang, P., J. Cutler, J. King and D. Palmer, 2004 Mutation of the regulator of G protein signaling Crg1 increases virulence in *Cryptococcus neoformans*. Eukaryot. Cell 3: 1028–1035.

Wang, Z., A. Wilson and J. Xu, 2015 Mitochondrial DNA inheritance in the human fungal pathogen *Cryptococcus gattii*. Fungal Genetics and Biology 75: 1–10.

Wilson, A. J., and J. Xu, 2012 Mitochondrial inheritance: diverse patterns and mechanisms with an emphasis on fungi. Mycology 3: 158–166.

Xu, J., 2005 The inheritance of organelle genes and genomes: patterns and mechanisms. Genome 48: 951–958.

Xu, J., R. Y. Ali, D. A. Gregory, D. Amick, S. E. Lambert et al., 2000 Uniparental mitochondrial transmission in sexual crosses in *Cryptococcus neoformans*. Current Microbiology 40: 269–273.

Yan, Z., C. M. Hull, J. Heitman, S. Sun and J. Xu, 2004 *SXI1*α controls uniparental mitochondrial inheritance in *Cryptococcus neoformans*. Current Biology 14: R743–744.

Yan, Z., C. M. Hull, S. Sun, J. Heitman and J. P. Xu, 2007a The mating-type specific homeodomain genes *SXI1*α and *SXI2*a coordinately control uniparental mitochondrial inheritance in *Cryptococcus neoformans*. Current Genetics 51: 187–195.

Yan, Z., Z. Li, L. Yan, Y. Yu, Y. Cheng et al., 2018 Deletion of the sex-determining gene *SXI1*α enhances the spread of mitochondrial introns in *Cryptococcus neoformans*. Mobile DNA 9: 24.

Yan, Z., S. Sun, M. Shahid and J. Xu, 2007b Environment factors can influence mitochondrial inheritance in the fungus *Cryptococcus neoformans*. Fungal Genetics and Biology 44: 315–322.

Yan, Z., and J. Xu, 2003 Mitochondria are inherited from the *MAT*a parent in crosses of the basidiomycete fungus *Cryptococcus neoformans*. Genetics 163: 1315–1325.

Zhou, Q., H. Li and D. Xue, 2011 Elimination of paternal mitochondria through the lysosomal degradation pathway in *C. elegans*. Cell Res 21: 1662–1669.

